# Cortical miR-709 links glutamatergic signaling to NREM sleep EEG slow waves in an activity-dependent manner

**DOI:** 10.1101/2022.09.24.508386

**Authors:** Konstantinos Kompotis, Géraldine M. Mang, Jeffrey Hubbard, Sonia Jimenez, Yann Emmenegger, Christos Polysopoulos, Charlotte N. Hor, Leonore Wigger, Sébastien S. Hébert, Valérie Mongrain, Paul Franken

**Affiliations:** Center for Integrative Genomics, University of Lausanne, Lausanne, VD, CH-1015, Switzerland; Genomic Technologies Facility, Center for Integrative Genomics, University of Lausanne, Lausanne, VD, CH-1015 Switzerland; Centre de recherche du CHU de Québec-Université Laval, CHUL, Axe Neurosciences, Québec, Canada; Faculté de médecine, Département de psychiatrie et de neurosciences, Université Laval, Québec, Canada; Center for Advanced Research in Sleep Medicine, Hôpital du Sacré-Coeur de Montréal, Montréal, QC H4J 1C5, Canada; Department of Neuroscience, Université de Montréal, Montréal, QC H3T 1J4, Canada; Department of Biostatistics, Epidemiology, Biostatistics and Prevention Institute (EBPI), University of Zurich, Zurich, ZH, CH-8057 Switzerland

**Keywords:** MicroRNAs, miR-709, *Dicer*, sleep deprivation, EEG delta power, glutamatergic receptor activity, endosomal trafficking, cortical plasticity

## Abstract

MicroRNAs (miRNAs) are key post-transcriptional regulators of gene expression that have been implicated in a plethora of neuronal processes. Nevertheless, their role in regulating brain activity in the context of sleep has so far received little attention. To test their involvement, we deleted mature miRNAs in post-mitotic neurons at two developmental ages, i.e., in early adulthood using conditional *Dicer* knockout (cKO) mice and in adult mice using an inducible conditional *Dicer* cKO (icKO) line. In both models, electroencephalographic (EEG) activity was affected and the response to sleep deprivation (SD), altered; while rapid-eye-movement sleep (REMS) rebound was compromised in both, EEG delta (1-4 Hz) power during non-REM sleep (NREMS) was reduced in cKO mice and increased in icKO mice. We subsequently investigated the effects of SD on the miRNA transcriptome and found that the expression of 48 forebrain miRNAs was affected, in particular, the activity-dependent miRNA miR-709. *In vivo* inhibition of miR-709 in the brain increased EEG power during NREMS in the slow-delta (0.75-1.75 Hz) range, particularly after periods of prolonged wakefulness. Transcriptome analysis of primary cortical neurons *in vitro* revealed that miR-709 regulates endosomal trafficking and glutamatergic receptor activity. A subset of the genes involved in glutamatergic transmission was affected also in the cortices of sleep-deprived, miR-709-inhibited mice. Our data implicate miRNAs in the regulation of EEG activity and indicate that miR-709 links neuronal excitability during wakefulness to brain synchrony during sleep, likely through the regulation of endosomal trafficking and glutamatergic signaling.

**Significance Statement:** MicroRNAs (miRNAs) are key regulators of gene expression playing vital roles both in postnatal brain development and its functioning in adult organisms. Here, we highlight a fundamental role for miRNAs in shaping EEG slow waves, which reflect synchronous neuronal firing, characteristic of NREM sleep (NREMS) in the adult murine cortex. Disruption of the miRNA-biogenesis machinery affected brain synchrony differently, depending on when it occurred during development. Moreover, sleep deprivation altered the expression of several miRNAs in a brain-region specific manner. Among those, we identified miR-709 to affect the expression of genes involved in endosomal-trafficking and glutamatergic-transmission, thereby linking neuronal activity during wakefulness to slow EEG waves during subsequent sleep. The current study causally implicates this specific miRNA and the molecular pathways it targets in modifying the generation of NREMS EEG slow waves, which are important in synaptic plasticity and brain functioning.

## Introduction

The molecular basis of sleep regulation has been under the scope of researchers for at least 20 years, in an effort to better understand sleep itself (1–4), reviewed in (5) It has become evident across species that after periods of extended wakefulness, either spontaneous or experimentally enforced (i.e., sleep deprivation or SD), brain gene expression is altered to accommodate neuronal responses in excitability and synaptic plasticity during sleep ((6), reviewed in (7)). These molecular changes are sufficient to modify global brain activity, as assessed by electroencephalographic (EEG) recordings (8, 9). Molecular players involved in sleep include immediate early genes, adhesion and scaffolding proteins, ion channels, synapse-related genes, neurotransmitters, protein transporters, genes involved in lipid and energy metabolism (reviewed in (3)), and more recently, in endosomal trafficking (10) and autophagic processes (11, 12). Although an important part of the molecular substrate of sleep has been unveiled, the involvement of gene expression regulatory mechanisms, other than transcription factors, has yet to be fully understood. One such mechanism is the fine-tuning of gene expression at the post-transcriptional level by microRNAs (miRNAs).

MiRNAs are ~22 nucleotide short, non-coding RNAs that must undergo a maturation process requiring the enzymes DROSHA/DGCR8 and DICER (11). Mature miRNAs exert their function by guiding the RNA-induced silencing complex (RISC) to their mRNA targets (13). Once bound to their target, miRNAs either destabilize the targeted transcript leading to repressed translation, or induce degradation (14). They have been implicated in a plethora of physiological and pathological processes, such as neuronal development and neurodegeneration (15), as well as modifying neuronal excitability via the regulation of the expression of genes encoding ion-channel (16), cytoskeletal (17), and scaffolding-protein (18), amongst others. Despite their known role as central transcriptional regulators in all molecular pathways studied, only a limited number of studies in rats (19–21), humans (22), and flies (23, 24) have investigated their involvement in the regulation of sleep. We previously identified 10 miRNAs among the genes most consistently affected by SD (1). More recently, miR-137 was found to modulate wakefulness in mice (25). Taken together, these observations suggest that miRNAs represent an additional, evolutionarily conserved, layer of gene regulation implicated in sleep processes. Nevertheless, their roles in the regulation of brain synchrony and associated processes during sleep have not been assessed, and their key downstream pathways in the context of sleep remain unidentified.

In the current study, we show that miRNAs regulate EEG brain activity across sleep-wake states in the adult mouse. In two conditional *Dicer* knockout mouse lines, each providing a different temporal blockade of miRNA maturation in forebrain excitatory neurons, REM sleep (REMS) was compromised, and EEG activity after SD was altered. We identified SD-induced changes in miRNA expression in the brain and subsequently inhibited *in vivo* the top differentially expressed miRNA, miR-709. Inhibition of miR-709 *in vivo* increased the number of slow waves in the slow-delta (0.75-1.75 Hz) band, while its inhibition in primary cortical neuronal cultures revealed genes involved in endosomal trafficking and glutamatergic receptor signaling being most affected. The *in vitro* differentially regulated genes were also assessed *in vivo* in miR-709 inhibited animals, where the cortical expression of genes related to glutamatergic synaptic transmission was again affected the most. Our observations support the involvement of miRNAs in the regulation of cortical slow-wave characteristics and implicate miR-709 in the control of pathways directly linked to neuronal synaptic plasticity (26–29)

## Results

### Constitutive conditional *Dicer* deletion impacts EEG spectral power, and abolishes NREMS EEG and REMS amount in response to sleep deprivation

To establish a role for mature miRNAs in sleep regulation in the adult mouse, we assessed the consequences of *Dicer* ablation in post-mitotic excitatory neurons. First, 12-weeks-old conditional *Camk2a-Dicer^fl/fl^* knockout male mice (cKO, n=8, referred to as *Dicer^fl/fl^)* lacking forebrain *Dicer* from 1.5 months of age were contrasted to their control littermates (n=6). cKO mice exhibited a highly aberrant low voltage EEG signal (**Figure S1A**), which, however, did not impede sleep-wake state identification. Absolute levels of EEG spectral power were reduced in cKO mice in all sleep-wake states and EEG frequencies >2.0 Hz compared to the *Dicer^fl/+^* littermate controls (**Figure S1B**). Despite these differences in EEG signal amplitude, the typical sleep-wake driven dynamics in NREMS EEG delta (1-4 Hz) power (30), appeared largely preserved. Nevertheless, differences in the range of these relative changes were observed. At light onset, the beginning of the main rest phase in baseline, cKO mice showed significantly higher EEG delta power compared to the control group (168% vs. 155%, **Figure 1A**, *top panel;* both were expressed relative to levels reached in the last 4h of baseline light period when delta power no longer declines further (30)). cKO mice slept less during the hour before and after light onset than their control littermates, and also lacked the additional resting period during the second half of the dark period (‘nap’ at ZT20-22; **Figure 1A**, *mid panel*), typical in this strain of mice (31, 32). These differences in the baseline sleep-wake distribution might have contributed to a higher sleep homeostatic drive and associated higher relative EEG delta levels observed in cKO mice at the beginning of light period. After 6 hours of SD, control mice exhibited the expected surge of NREMS EEG delta power (31)reaching 230% over the baseline reference (**Figure 1A**, *top panel*). Conversely, EEG delta power in *Dicer* cKO mice did not significantly increase over levels reached at light onset in baseline (183% of reference) when the SD was initiated, and levels reached after SD were lower than in control mice.

**Figure 1.**
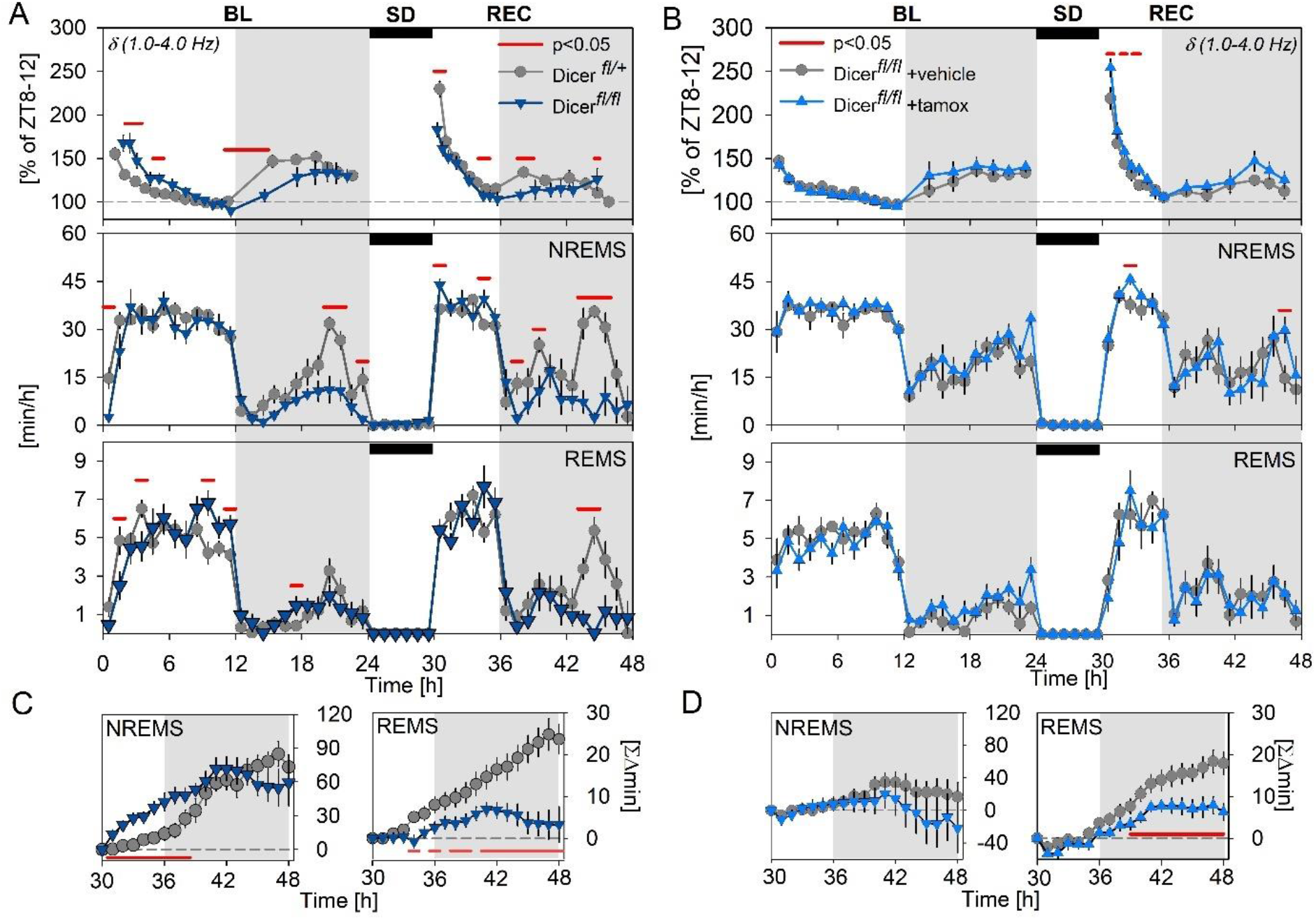
Temporal control of Dicer deletion defines its impact on EEG delta power. Sleep parameters were assessed in conditional (cKO, A and C) and inducible-conditional (icKO, B and D) Dicer knockout mice. **A and B.** From top to bottom: Mean time course of NREMS EEG delta power (δ referenced to baseline ZT8-12 EEG delta power) and NREMS and REMS time, during 24h baseline (BL) and 18h of recovery (REC) from a 6h sleep deprivation (SD; black rectangles) starting at light onset (ZT0-6, dashed grey background). Grey areas delineate the dark periods. Significant changes for EEG delta power, NREMS, and REMS, are depicted by red horizontal lines above datasets (2-way rANOVA interaction factors genotype x time p < 0.001; post-hoc t-tests p < 0.05). **C and D.** NREMS (left panel) and REMS (right panel) rebound for the 18h of recovery after SD as accumulated differences from corresponding baseline hours. Significantly smaller rebounds in sleep time are marked by the red horizontal lines underneath datasets (t-tests p<0.05). N =*Dicer^fl/+^:* 8*; Dicer^fl/fl^:* 6; *Dicer^fl/fl^* + vehicle: 6; *Dicer^fl/fl^* + tamoxifen: 7.

Except for the dark period time points mentioned above (Zeitgeber Time ZT20-22 and ZT23-1), time spent in NREMS in baseline did not differ between cKO mice and controls (**Figure 1A**, *middle panel*). Also, during the dark period following the SD, cKO mice spent less time in NREMS, especially between ZT19 and −22 (**Figure 1A**, *middle panel).* During the dark period, REMS was similarly reduced in cKO, although significantly so only following SD (**Figure 1A**, *bottom panel*). Although cKO mice initially spent more time in NREMS during the 6 hours after SD, compared to corresponding baseline hours, this difference in accumulated extra NREMS time dissipated over the course of the subsequent dark period (**Figure 1C,** *right panel*). In contrast, while control mice did gain extra REMS during the recovery period, SD in cKO mice failed to induce a compensatory response (+23.6 vs. +3.3 min, respectively, by the end of the recovery dark period; **Figure 1C,** *right panel*).

Given the genotype differences in the dynamics of EEG delta power, we next investigated the effect of conditional *Dicer* deletion on EEG slow-wave (SW) features during NREMS in the time-domain, such as SW density and amplitude. We selected temporal windows reflecting different levels of sleep pressure; i.e. the end of the baseline rest phase, after the main waking period in the baseline dark period, as well as the period directly following SD. Both SW density and amplitude appeared reduced in cKO mice, regardless of sleep pressure, except for the density of the slowest SWs (<1.25Hz; **Figures 2A** and **B**). Consistent with our previous observations (33), two distinct SW sub-bands were evident, each responding differently to extended wakefulness. We thus analysed the delta power dynamics separately for slower (δ1: 0.5-1.75 Hz) and faster (δ2: 2.0-3.5 Hz) delta frequencies. As shown in **Figure 2E**, the power of both the δ1 *(upper panel)* and δ2 *(bottom panel)* sub-band was higher during the first hours of light period, and lower after sleep loss in cKO mice, compared to mice with functional *Dicer*. However, the differences during the dark period were present solely in the δ1 band, which in contrast to δ2, was reduced in the cKO group.

**Figure 2:**
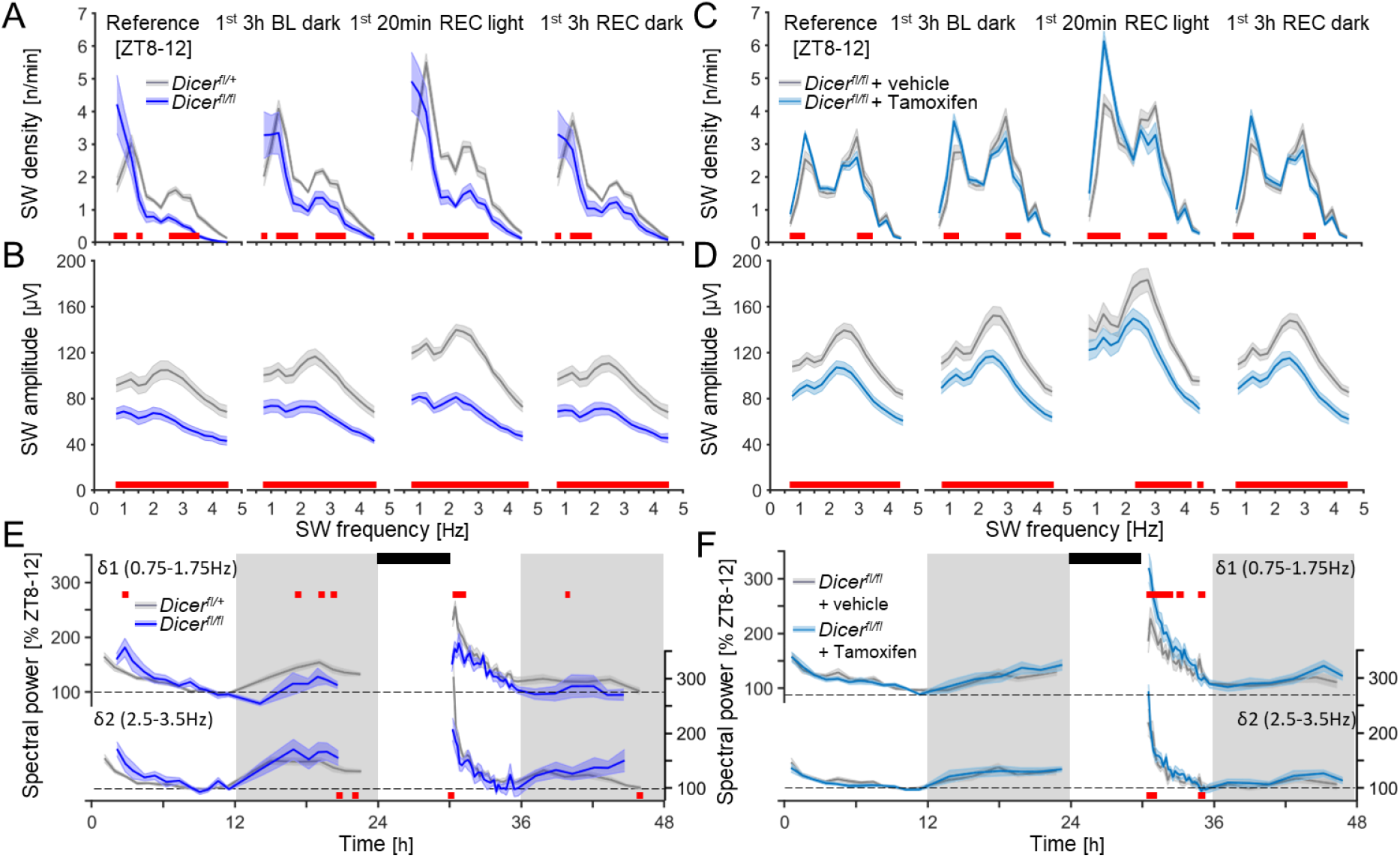
Dicer deletion modifies EEG slow-wave (SW) properties following spontaneous and enforced (sleep deprivation) waking. NREMS EEG SW parameters were assessed in conditional (A, B, and E; cKO, dark blue) and inducible-conditional (C, D, and F; icKO, light blue) *Dicer* knockout mice. **A and C.** SW density in cKO and icKO mice at different times of the experiment during which sleep pressure is assumed to differ; i.e., the reference period (ZT8-12) with lowest sleep pressure in baseline (BL), first 3h of the dark period (ZT12-15) with highest BL sleep pressure, the first 20min of recovery (REC) from SD with highest sleep pressure during the experiment, and the first 3h of the REC dark period, when sleep pressure is lower than in BL as mice usually sleep more (31). SW frequencies with significant changes in density, compared to controls (grey), are depicted by the red squares (2-way rANOVA interaction factors genotype × frequency; cKO: p < 0.001 for all periods; icKO: p < 0.001 for all periods; post-hoc Tukey: p<0.05). SW density is frequency corrected (n/frequency), to better illustrate SW bimodality. **B and D**. SW amplitudes in cKO and iCKO for the same periods as in A and C. Changes from control (grey) are depicted with red squares (2-way rANOVA interaction factors genotype × frequency; cKO: p < 0.001 for all periods; icKO: p < 9E-5 for all periods). **E and F.** Time-course of δ1 (0.75-1.75 Hz; **top**) and δ2 (2.5-3.5 Hz; **bottom**) spectral power during baseline and following 6h SD (black rectangles), in *Dicer* knockout mice and controls. Statistics (2-way rANOVA interaction genotype × time; cKO: p < 0.001 for both δ1 and δ2; icKO: δ1: p <0.001; δ2: p = 0.002). All values represent means (solid lines) ± SEM (shaded areas). Power was expressed as % of mean δ1 and δ2 power, respectively, during the last 4h of the light period of baseline (ZT8-12). N =*Dicer^fl/+^*: 8; *Dicer^fl/fl^*: 6; *Dicer^fl/fl^* + vehicle: 6; *Dicer^fl/fl^* + tamoxifen: 7.

The aberrant EEG signal in cKO mice supports an essential role for miRNAs in brain development and function, consistent with previous literature (34, 35). Sleep-wake driven changes in brain activity of these mice, as observed by their EEG SW dynamics, might reflect deficits in neuronal synchrony. Given that previous research has reported cortical neuroinflammation starting at 2.5 months and neurodegeneration at 6 months of age for this particular mouse line (29, 30), neuronal synchrony deficits could have emerged from impaired neuronal communication, potentially due to the onset of neuroinflammatory processes during early adulthood in the absence of functional *Dicer.* Thus, we next chose to explore miRNA regulation of sleep EEG activity using a more temporally regulated approach.

### Inducible *Dicer* deletion in excitatory neurons of the adult brain amplifies the increase in EEG delta power and reduces the REMS rebound after sleep deprivation

To determine the consequences of acute neuronal miRNA depletion in the post-mitotic neurons of adult mice, sleep and EEG features of inducible conditional *Dicer* knockout (icKO; *Dicer^fl/fl^* + tamoxifen; 12 weeks old, n=7) male mice were assessed and compared to their non-induced control littermates (*Dicer^fl/fl^* + vehicle, n=6). We previously confirmed that *Dicer* deletion occurred ~3 weeks following tamoxifen injections in 2-month-old mice (see (36)). As in cKO mice, we observed a reduction in the absolute levels of EEG spectral power in the icKO mice in all 3 sleep-wake states, particularly in the range of 2.0-10.0 Hz for wakefulness and 2.0-30.0 Hz for NREMS and REMS (**Figure S1D**). Nevertheless, the relative spectral profiles in icKO seemed relatively unperturbed compared to those obtained in cKO mice. NREMS EEG delta power dynamics did not differ between the icKO and control groups under baseline conditions but, contrary to cKO mice, SD in icKO mice resulted in a significantly larger increase compared to controls (**Figure 1B**, upper panel). Major differences in sleep-wake state distribution were not observed, with the exception of a reduced REMS rebound post SD in icKO mice (**Figure 1C**, *right panel*).

Additionally, icKO mice exhibited higher SW density in the δ1 range (0.75-1.75 Hz), particularly after SD (1^st^ 20min REC), while faster SWs (δ2; 2.5-3.5 Hz) were less prevalent (**Figure 2C**). Conversely, SW amplitude was reduced in icKO mice at all periods, regardless of sleep pressure (**Figure 2D**). Note that the amplitude of SWs in the δ1 range reached levels similar as the control group post SD (**Figure 2D,** 1^st^ 20min REC), while that of SWs in the δ2 range remained lower. The larger post-SD increase in NREMS EEG power in the delta (1-4 Hz) range in icKO mice (**Figure 1B**), was thus largely due to the differences in the lower δ1 sub-band activity (**Figure 2F**).

Together, our observations in both *Dicer* knockout mouse models suggest that interfering with miRNA maturation in excitatory neurons yields broad changes in brain activity during both wakefulness and sleep. Specific to NREMS SWs, amplitude was reduced while incidence of slower SWs was increased and that of faster SWs decreased in both lines. The two lines also shared a reduced REMS rebound after SD while the delta rebound showed opposite effects attributable in large part to differences in the δ1 sub-band which is reported to respond less to sleep pressure (33).

### Sleep deprivation increases miR-709 expression in the forebrain

To identify specific miRNAs affected by enforced waking, we next compared miRNA expression after SD in murine brains. Among the 611 miRNAs quantified, 48 miRNAs were differentially expressed in sleep-deprived forebrains (27 up-, 21 downregulated, moderate t-statistic p<0.05), but only the increase in miR-709 levels survived correction for multiple testing (1.3-fold; FDR<0.05; **Figure 3A**). Of note, multiple miRNAs may target the same gene(s) and thus potentially the same pathways (37). To identify these pathways, we investigated the network of shared validated targets from the 48 differentially regulated forebrain miRNAs using the miRNET resource ((38, 39), **Figure 3B**). Amongst the pathways that were significantly overrepresented in the set of shared target genes were *‘axon guidance’* (KEGG: mmu04360), *‘regulation of actin cytoskeleton’* (KEGG: mmu04810), *‘regulation of intracellular transport’* (GO: 0006886), *‘endosomal transport’* (GO: 0016197) and *‘early endosome’* (GO: 0005769).

**Figure 3.**
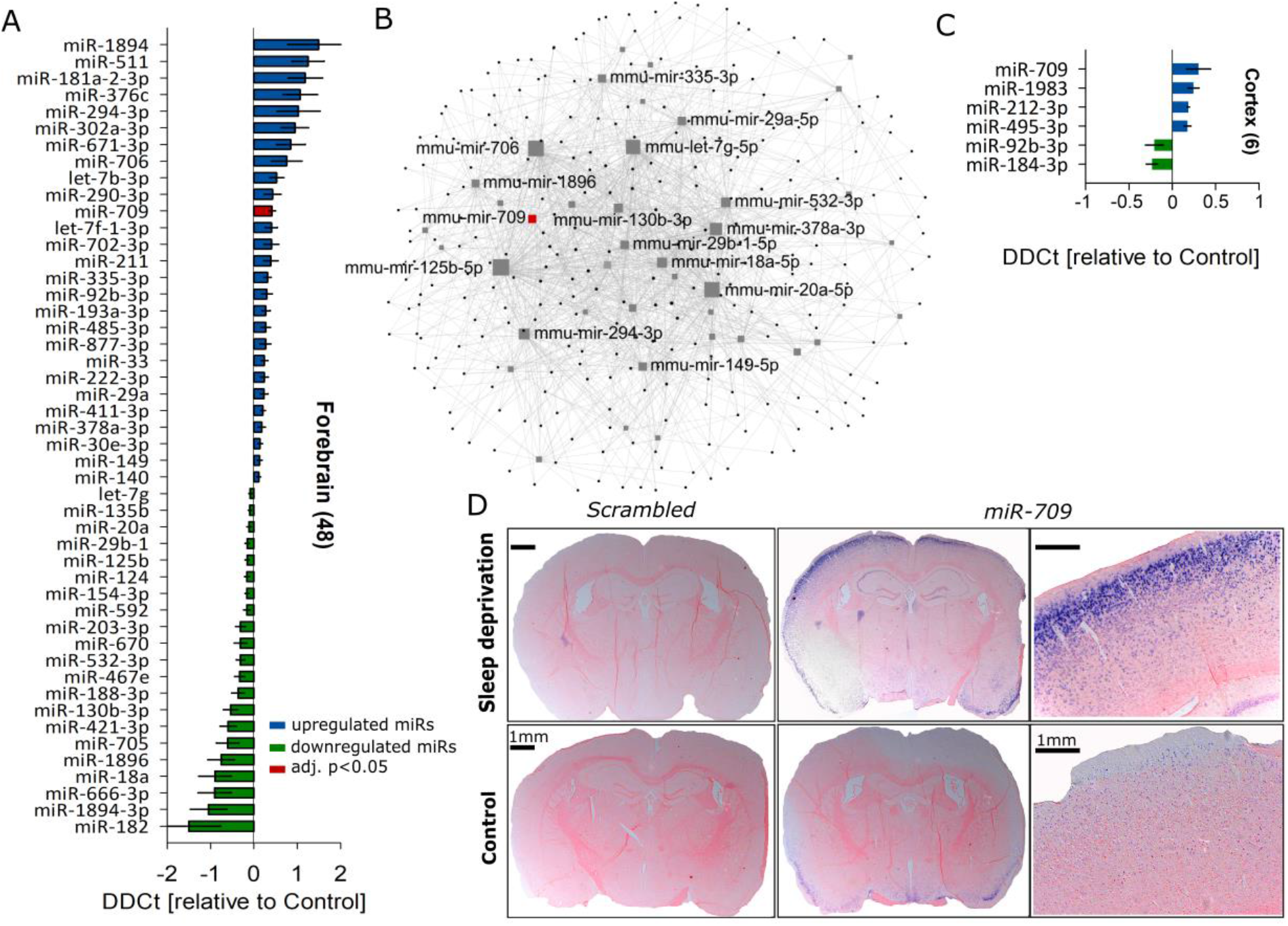
miR-709 is the top differentially regulated miRNA after sleep deprivation. Bar charts represent the relative expression levels of significantly altered miRNAs between 6h-sleep deprived (SD, n=5) and control (Ctrl, n=5) animals. Error bars show the pooled SEM of SD and Ctrl samples. **A.** SD significantly affected the forebrain expression of 48 (out of 611 tested miRNAs) of which 27 were up-(blue) and 21 downregulated (green; SD vs Ctrl; moderate t-statistic, p<0.05). Note that only miR-709 expression (red) survived multiple testing correction (FDR<0.05). **B.** Network of all differentially regulated miRNAs in the forebrain after SD and their validated targets (black dots; miRTarBase v8.0 and TarBase 8.0, see **Supp. Table S1** for the complete miRNA-target list), showing the names of the top 15 interconnected forebrain miRNAs (grey squares), including miR-709 (red square). Note that larger squares are more interconnected. Only the shortest path between two hub nodes is visualised (40 miRNA nodes; 273 gene nodes; 1421 edges; for participating pathways see **Supp. Table S2**). **C.** Mean differential miRNA expression in cortex after 6h SD, where 4 miRNAs increased and 2 decreased their expression (SD vs Ctrl; moderate t-statistic, p<0.05, FDR>0.05). For hippocampal miR expression, see **Figure S3E**. **D.** *In situ* hybridization of two representative sleep-deprived (top panels) and control (bottom panels) mice, with either a miR-709-specific probe (four right panels) or a scrambled miRNA probe (two left panels). *In situ* hybridization confirms that SD increased miR-709 expression in the cortex. No signal was detected with the scrambled probe.

Since miRNA expression and function have been found to be brain region-specific (40, 41), we next investigated cortices (**Figure 3C**) and hippocampi (**Figure S3E**) of sleep deprived mice in a second, independent cohort. Simple t-tests suggested a significant SD-driven upregulation of miR-709 also in the cortex (1.2-fold; p=0.03), nevertheless miRNA expression changes in both tissues failed to reach statistical significance after correction for multiple testing (FDR > 0.05; **Figure 3C**). I*n situ* hybridization confirmed the SD-induced increase in miR-709 cortical levels. This increase was most pronounced in cortical layers 2 and 3, encompassing at least the somatosensory cortex (**Figure 3D**). In summary, miR-709 levels were significantly increased by sleep loss in the murine forebrain. We thus further investigated the role of this miRNA in the wake-dependent changes in brain activity during sleep.

### *In vivo* miR-709 inhibition increases the prevalence of NREMS δ1 SWs, particularly after periods of prolonged spontaneous wakefulness

As SD upregulated miR-709, we hypothesized that *in vivo* miR-709 inhibition would reveal its role in regulating the changes in NREMS SWs after prolonged periods of wakefulness. We therefore injected 12-week-old male mice intracerebroventricularly (ICV) with miR-709-specific locked-nucleic-acid (LNA) inhibitors (i-miR-709, n=7), non-specific scrambled probes (scram, n=7), or vehicle (artificial cerebrospinal fluid; aCSF, n=5), forty-eight hours prior to the onset of EEG recordings (see Methods). Inhibition of miR-709 in vivo resulted in a consistently higher SW density restricted to the lower delta frequencies (δ1; 0.75-1.75 Hz; **Figure 4A**, upper panel), while SW amplitudes remained unaffected (**Figure 4A**, lower panel). Subsequent analysis of NREMS EEG delta power dynamics showed that δ1 power was higher in NREMS episodes after spontaneous wakefulness during the baseline dark period but did not reach statistical significance after SD (**Figure 4B** and **S2A**). No robust differences were overall observed in the time spent in NREMS and REMS between the three groups (**Figures 4C** and **D**). Moreover, the NREMS and REMS rebounds after SD were unaffected (**Figure 4D**).

**Figure 4:**
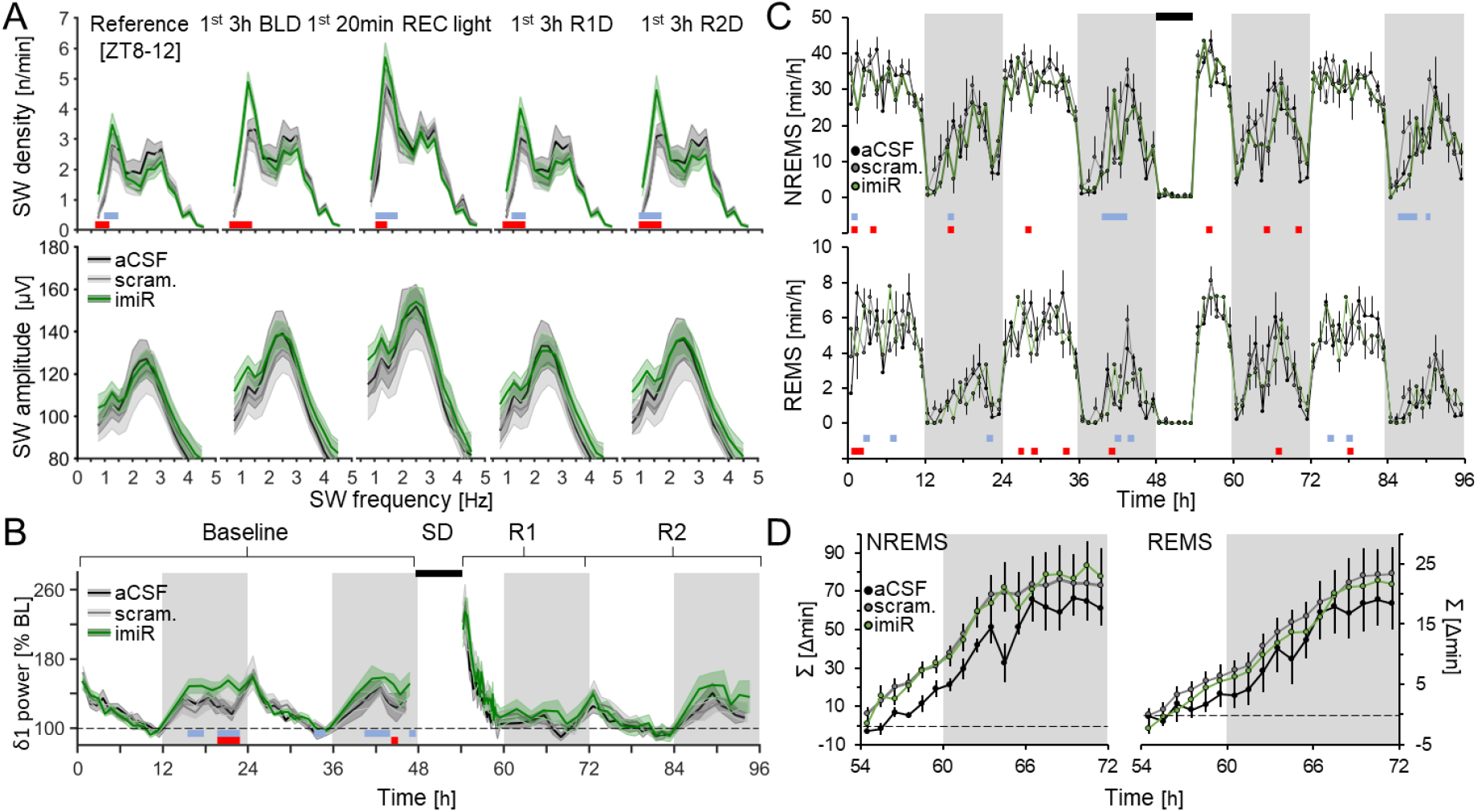
miR-709 inhibition increased SW density in the δ1 range during NREMS. **A.** Mice inhibited for miR-709 (imiR, **green**) show significant increases in the density (**top**) of δ1 SWs following spontaneous and enforced waking, compared to either scrambled (**grey**) or aCSF (**black**) controls during (from left to right): the last 4 hours of light period (Reference [ZT8-12]), the first 3 hours of baseline dark period NREMS (1^st^ 3h BLD), the first 20 minutes of NREMS after sleep deprivation (1^st^ 20min REC light), and the first 3 hours during the two subsequent recovery dark periods (1^st^ 3h R1D and R2D, respectively). Red squares represent significant changes between aCSF and imiR, and blue between scrambled and imiR, groups (2-way rANOVA interaction factors genotype × frequency; p < 0.001 for all periods; post-hoc t-tests p<0.05). SW density is frequency corrected (n/frequency) to better illustrate SW bimodality. **B.** Time-course of NREMS δ1 (0.75-1.75 Hz) power during 48h of baseline and 42h following a 6h SD; i.e. recovery days R1 and R2. imiR mice had significantly higher δ1 power following spontaneous but not enforced waking (2-way rANOVA factors genotype × time; p = 0.04). Values are expressed as % of mean δ1 power during the last 4h of the light period of baseline (ZT8-12). Note that 2 baseline days were averaged for statistics. **C.** Time-course of NREMS (top) and REMS (bottom) amounts per hour (2-way rANOVA interaction factors genotype × time; NREMS: p = 0.006; REMS: p = 0.03; post-hoc tests <0.05). **D.** Accumulated differences in minutes between baseline and following 6h SD, in NREMS and REMS. All values represent means (solid lines) ± 1 SEM (shaded areas or bars). N = aCSF: 5; scrambled: 7; imiR: 7.

Taken together, these observations suggest a role for the mature miR-709 specifically in the regulation of NREMS EEG δ1 waves, which may also contribute to the similar δ1 density phenotype observed in the cKO and icKO mice (**Figure 2A and C**).

### miR-709 inhibition downregulates glutamatergic receptor genes and upregulates endosomal trafficking genes in primary cortical neurons

To reveal potential molecular pathways by which miR-709 inhibition affected activity of δ1 SWs, we next aimed at identifying its transcriptional targets. First, we quantified transcriptomic changes after inhibiting miR-709 in an *in vitro* assay, where inhibitors can be directly applied to the neuronal population of interest. Since miR-709 expression was previously observed in cortical neurons (42), its levels were affected by SD mainly in the cortex (**Figure 3C** and **D**), and a cortical origin has previously been suggested for slow SWs (33, 43), we treated cultured primary cortical neurons with either miR-709-specific LNA inhibitors (i-miR-709, n=6), scrambled LNA inhibitors (scr, n=6), or vehicle (ddH2O, n=6). RNA sequencing (RNA-seq) after treatment identified a total of 13’760 expressed genes. To assess the effect of miR-709 inhibition at the individual gene level, gene expression in the inhibitor-treated mice was contrasted to those of the scrambled-treated mice. We found 111 genes to be differentially expressed (FDR<0.05), with only six genes *(Akap12, s100a10, Acadm, Glipr2, Prrg3*, and *Hexb*) being upregulated by miR-709 inhibition (**Figure 5A**) and all others downregulated, including several ion channel encoding genes. Of note, considering changes in the transcriptome as a whole, and assessing the effect of miR-709 inhibition on networks of co-expressed genes acting together in the same pathway(s), is likely to provide a more complete picture on its regulatory role. Thus, Weighted Gene Co-expression Network Analysis (WGCNA) was used on all sequenced transcripts of all 18 samples from the three experimental groups to screen for gene sets and related pathways. The analysis revealed eleven clusters, or modules, each containing a variable number of genes (from 100 to 2000; **Figures S4A-C**). Four modules were found to significantly correlate with miR-709 inhibition (**Figure 5B**; *‘green’ r*=-0.97; p=9e-11; *‘purple’* r=0.84, p=1e-05; *‘black’* r=0.52, p=0.03; and *‘pink’* r=0.54, p=0.02). The most representative genes of each of these modules, i.e., the ‘hub genes’ identified by WGCNA (**Supp. Table S4**), are co-expressed and often interact biologically at the protein level (44), as illustrated by the protein-protein interaction (PPI) networks (**Figures 5C-D** and **S5**). Interestingly, in the hub genes from two *(‘black’* and *‘pink’*) out of the three positively correlated networks, there was a strong presence of Rab GTPases and genes directly involved in endosomal trafficking (**Figures 5C** and **S5B**; also see **Supp. Table S4**). Conversely, the negatively correlated network (‘*green*’), was heavily populated by ion-channel and synaptic-scaffold associated proteins, while it encompassed most of the 111 differentially expressed genes, with some (e.g. *Camk2g, Nreg3, Tbce*) emerging as hub genes. The topmost molecular functions (**Figure 5E**; *top panel*) were associated with *extracellular glutamate gated ion channel activity* (GO: 0005234; Strength: 1.29) and *ionotropic glutamate receptor activity* (GO: 0004970; Strength: 1.27). Moreover, the main participating biological processes (**Figure 5E**; *bottom panel*) were *regulation of vesicle docking* (GO: 0106020; Strength: 1.72), *positive regulation of glutamate secretion, neurotransmission* (GO: 1903296; Strength: 1.72) and *positive regulation of long-term synaptic depression* (GO: 1900454; Strength: 1.6).

**Figure 5.**
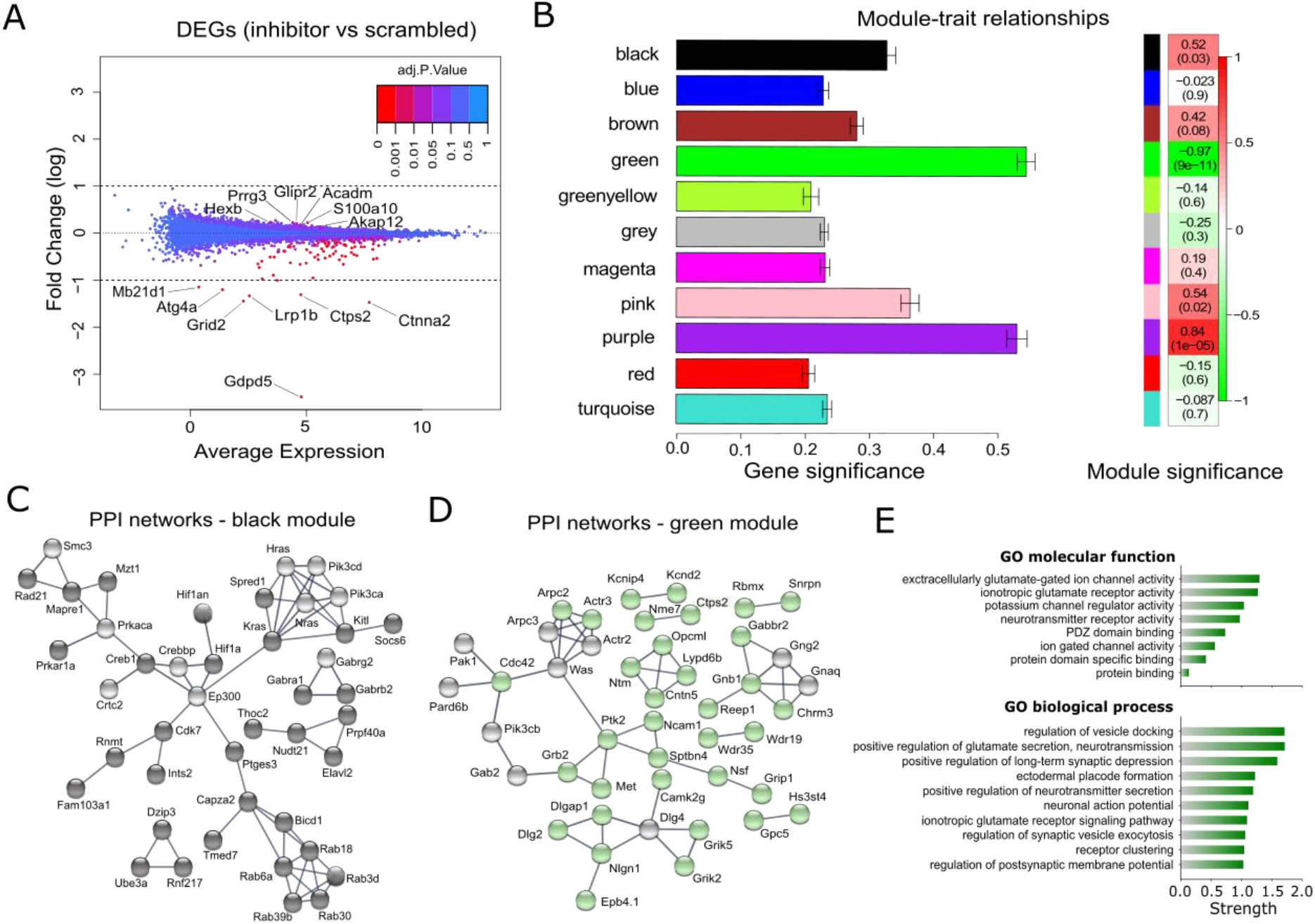
Differential expression and WGCNA analysis of genes affected by miR-709 inhibition in primary cortical cultures. **A.** Multivariate-analysis plot for the differential transcriptome analysis by RNA sequencing between the inhibitor- and scramble-treated groups. Dots represent the differential expression of a gene (DEG); 111 genes were significantly altered by miR-709 inhibition (FDR<0.05) some of which are named (for the full list, see **Supp. Table S3**). **B-E.** Weighted Gene Correlation Network Analysis (WGCNA) for all genes identified by RNA sequencing (see Methods and **Figure S4**). **B.** Gene and module significance calculation, given by the Pearson correlation of the corresponding module eigengene to the miR-709 inhibition. The table contains the coefficient (p-value) for each module-trait correlation, is color-coded according to the correlation coefficient (green to red), and is accompanied on the left by the corresponding module color. **C and D.** Visualization of the protein-protein interaction (PPI) networks of top interconnected (hub) genes of the black and green module (colored spheres) in STRING (45). Grey spheres represent genes directly linked to the green hub genes (first shell interactors; ‘*no more than 10’*option), which however were not included in our initial dataset. **E.** Top functional enrichment annotations of the green module hub genes for Gene Ontology (GO) molecular function (8) and biological process (top 10). Pathway terms are hierarchically ordered, based on their Strength [log10 (observed/expected) number of genes in a given pathway].

### *In vivo* miR-709 inhibition negatively impacts expression of ion transport genes

We subsequently assessed the levels of the 111 *in vitro* differentially expressed transcripts in the cortex of 12-weeks-old sleep-deprived male mice, ICV-injected with either miR-709-specific (i-miR-709, n=4) or scrambled (scr, n=5) LNA inhibitors (**Figures 6A** and **S6A-D**), same as for the EEG recordings. Inhibiting miR-709 significantly altered the expression of 23 of the 111 genes (19 up- and 4 downregulated), with *Farp1* and *Ctps2* being most affected (FDR<0.05; **Figure 6A**). Between the *in vitro* and *in vivo* experiments (**Figure 6B**), four genes were found commonly upregulated (*S100a10, Acadm, Hexb,* and *Glipr2*), and four commonly downregulated (*Grin1, Camk2g, Nrg3*, and *Tbce*). Additionally, PCA biplot analysis of expression levels of these eight genes successfully clustered them to the correct treatment group in all *in vivo* samples suggesting that they were co-regulated (**Figure S6D**).

**Figure 6.**
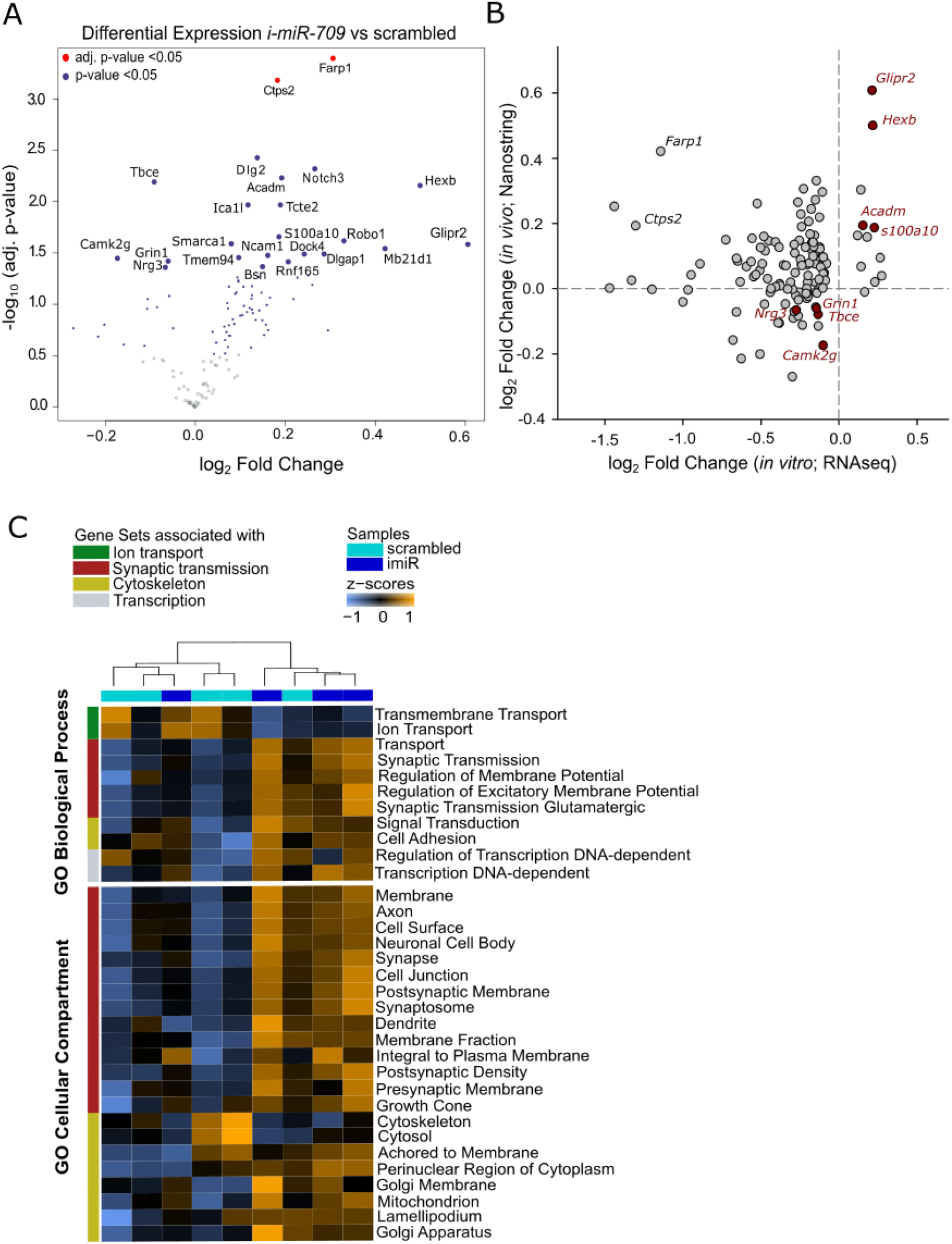
Differential gene expression after sleep loss in the cortex of i-miR-709 treated animals. Expression levels of the 111 differentially regulated genes from the *in vitro* miR-709 inhibition experiment were assessed in the cortical tissue of i-miR-709-(n=4) or scrambled-(n=5) injected animals after 6h SD. **A.** Volcano plot of differential gene expression between the two groups, displaying the log2 fold-change and the respective −log10 p-value for each individual gene. Genes with significantly altered expression (inhibited vs scrambled) are depicted by the blue (t-test, p-value<0.05) and red (Benjamini-Hochberg, FDR<0.05) dots. **B.** Correlation between expression levels of the top 111 genes in the *in vitro* and *in vivo* miR-709 inhibition experiment. Note that four genes are found to be commonly up- and four downregulated in both datasets. **C.** Heatmap of gene sets and their contributing Gene Ontology (GO) pathways in each sample. Out of the 48 GO pathways identified, only those related to GO Biological Process (11) and GO Cellular Compartment (22) were visualized. Samples are clustered in the dendrogram according to their z-transformed pathway scores (see Methods). GO annotations are shown on the right side of the heatmap.

We then investigated the participating pathways of gene sets affected by the *in vivo* miR-709 inhibition. All genes quantified were first clustered into gene sets (see Methods), based on the Gene Ontology (GO) annotations defined for each gene in the Gene Set Knowledge Base (GSKB; (46)). Gene set analysis (GSA) revealed that the expression levels of genes associated with *ion transport* decreased in murine cortices when miR-709 was inhibited, while genes involved in *glutamatergic synaptic transmission* showed increased expression (**Figure 6C**). Taken together with the pathways identified for the *‘green’* hub genes (**Figure 5D**), our observations pinpoint the *ionotropic receptor activity* and *glutamatergic neurotransmission* as pathways consistently targeted by miR-709 both in cortical tissue under sleep deprived conditions and in cultured cortical neurons.

## Discussion

In this study, we show that ablation of mature miRNAs in two different *Dicer* knockout models led to alterations in brain synchrony, particularly in response to sleep loss. We identified miRNAs affected by sleep loss in the mouse forebrain and provided evidence that cortical miR-709 regulates EEG slow waves during NREMS. Using *in vitro* and *in vivo* assays, we argue that miR-709 exerts its effects through post-transcriptional regulation of pathways essential for activity-dependent synaptic plasticity and neuronal excitability. Our findings establish that forebrain miRNAs regulate brain synchrony during NREMS in the mouse.

### Ablation of the micro-RNA maturation machinery in excitatory neurons dysregulates EEG responses to sleep loss

Notably, early knockout of *Dicer* in other mouse models has been reported to be deleterious for brain development, with overt signs of neurodegeneration and severe neuroinflammation at 2.5 months of age, resulting in early adulthood death (47, 48). In the current study, cKO mice were recorded at 2.5-3 months of age, when neuroinflammatory processes set in, and thus might underlie the highly aberrant EEG signals observed in all sleep-wake states (**Figures 2** and **S1**). Indeed, inflammation has been previously reported to alter brain oscillatory function regardless of sleep-wake state (49, 50). Considering a neuroinflammatory phenotype, it might seem surprising that sleep-wake states appeared little affected; the time spent asleep and the amplitude of the diurnal sleep-wake distribution of cKO mice were by and large normal. Moreover, the relative changes in NREMS EEG delta power still followed the sleep-wake distribution although lower levels were reached after extended periods of spontaneous and enforced wakefulness. Whether these blunted delta power dynamics point to a slower build-up of the sleep homeostatic process that it is thought to reflect (33, 51) or relates to abnormally synchronized brain activity (49) is difficult to discern in the context of our study. Nevertheless, the profound and constitutive reduction in both the density and amplitude of EEG SWs during NREMS we observed in cKO mice seems to support the latter.

Although icKO mice have a normal development, they are not without issues, as others have reported signs of neurodegeneration in specific brain regions several weeks after tamoxifen injection (52, 53). Moreover, we previously observed a severe, but transient, metabolic disruption after tamoxifen injection in *Dicer* icKO mice, while brain transcriptome analysis in cortex and hippocampus pointed to increased neuronal excitability at 4 and 8 weeks after functional *Dicer* ablation (36). Indeed, loss of *Dicer* in adult forebrain neurons has been demonstrated to affect plasticity and increase neuronal excitability, at least in CA1 pyramidal neurons (54). As previously suggested, *Dicer* ablation increases neuronal excitation-dependent responsiveness and disrupts neuroprotective homeostatic mechanisms against overactivation (54). In our hands, they exhibited a larger increase in EEG delta power in response to enforced wakefulness. This might provide a further link to the paradoxical transient enhancement of memory that was initially observed after induced *Dicer* loss (52, 54). In the current study, deletion of *Dicer* in icKO mice resulted in broad EEG alterations, both in density and amplitude, in subgroups of NREMS slow waves, namely δ1 and δ2. On the other hand, miR-709 inhibition in cortical neurons exhibited a more band- and state-defined EEG phenotype, towards the same direction as icKO mice. Considering that the inducible *Dicer* deletion affected most mature miRNAs, it seems plausible that miR-709 is amongst the miRNAs contributing to the observed phenotype in the icKO model. Together, our observations in both *Dicer*-ablated mouse models suggest that mature miRNAs in excitatory forebrain neurons are necessary for brain synchrony, both under physiological conditions and after sleep loss, as well as for REM homeostasis.

### miR-709 inhibition upregulates expression of endosomal trafficking/autophagy genes, while genes linked to NMDAR-dependent signaling are downregulated

Validated targets of the differentially expressed miRNAs after SD in the forebrain, were found significantly enriched in genes associated with the regulation of cytoskeleton and endosomal transport. Additionally, miR-709, as well as miR-706, have been found to target transcripts of protein kinase C alpha (PRKCA) (55, 56), a suppressor of autophagy (57) and regulator of endosomal trafficking (58). Most importantly, miR-709-driven suppression of genes involved in cytoskeletal organization, as well as endosomal recycling and cell adhesion, was previously observed in non-neuronal tissue (41).

Since miRNAs supress gene expression, an inhibition of miR-709 levels can be expected to result in an upregulation of its direct targets. Thus, at least some of the shared upregulated genes in our datasets could be direct targets of miR-709. Between the *in vitro* and *in vivo* model of miR-709 inhibition in the current study, four up-*(Glipr2, S100a10, Hexb* and *Acadm*) and four down-regulated (*Tbce, Nrg3, Camk2g* and *Grin1*) transcripts were shared. Indeed, *Glipr2* (aka *Gapr-1*), a negative regulator of autophagy (59), is a predicted target of miR-709 (TargetScanMouse, v8.0, 2021). *S100a10* (aka *p11*), has been involved in autophagic pathways and controls the distribution of recycling endosomes (60), while hexosaminidase b *(Hexb)* is one of the main lysosomic enzymes for autophagy (61, 62). The strong representation of Rab GTPases in two out of the three upregulated gene networks after *in vitro* miR-709 inhibition (**Figure S4**), further suggests that endosomal trafficking/autophagic pathways were affected. The upregulated Rab transcripts (Rab3c/d, −6a/b, −18, −30, and −39b), encode core proteins of autophagy and endosomal trafficking in neurons (63–68), in certain cases specifically involved in NMDAR trafficking (67, 69). Interestingly, autophagic and endosomal trafficking processes have already been identified as essential for fast dynamics of NMDAR-dependent receptor trafficking and function at the synapse (70, 71). An aberrant endosomal/autophagic process alters synaptic neurotransmission (reviewed in (72)), thereby modifying the electrophysiological properties of the neuron (73), and the respective brain synchrony (74).

In agreement with these observations, the *in vitro* and *in vivo* shared downregulated genes after miR-709 inhibition were shown to be strongly involved in the electrophysiological properties of neurons, with the exception of *Tbce (tubulin-specific chaperone E)* essential for Golgi-linked cytoskeletal assembly, axonal tubulin routing (75), and Golgi SNARE-mediated vesicle fusion (75). Specifically, neuregulin-3 *(Nrg3)* regulates glutamatergic transmission via the SNARE complex (76), and excitatory inputs of cortical interneurons (77), while its dysregulation has been associated with hypofunction of the glutamatergic pathway in *Nrg3-*related mental illnesses (76). In general, neuregulins have been found to induce reduction of NMDA receptor currents, potentially by boosting the internalization of NR1, the fundamental subunit of NMDARs, via an actin-dependent mechanism (reviewed in (78)). Moreover, *Camk2g* (aka CaMKIIγ or γCaMKII) acts downstream of NMDAR signalling, to regulate experience-dependent transcription and memory (79). *Camk2g* is important for Ca^2+^/CaM shuttling to the nucleus thereby, triggering CREB phosphorylation and gene expression, a signalling pathway suggested to track and respond to neuronal depolarization rather than synaptic activity (80). Lastly, and most importantly, the downregulated *Grin1* is encoding the obligatory NR1 subunit of all NMDARs, thus delineating a common pathway between three out of the four downregulated genes. Combining findings from previous reports and our study, we suggest that miR-709 directly regulates the expression of genes implicated in endosomal trafficking and the autophagic machinery in cortical neurons, thereby impacting glutamatergic receptor trafficking and thus NMDAR-dependent neurotransmission.

### miR-709 inhibition increases NREMS slow-delta density potentially via transcriptional dysregulation of NMDAR-dependent receptor trafficking

The observed NREMS δ1 increase in the miR-709 inhibited mice persisted also after SD (**Figure 4A**), albeit not evident in the δ1 dynamics (**Figure 4B**). When comparing the δ1 SW density during the dark period and after SD for the miR-709-inhibited group, it becomes apparent that slow waves occurred in a similar rate after either spontaneous or enforced prolonged wakefulness. A potential mechanism for the impact of miR-709-inhibition on δ1 SW occurrence may lie with NMDAR-dependent signalling, since miR-709 was recently found to be upregulated after extrasynaptic-NMDAR-induced excitotoxicity (81). Previously, its expression was shown to increase in response to *in vivo* application of (S)-3,5-Dihydroxyphenylglycine (DHPG), an agonist of group I metabotropic glutamate receptors (82), as well as stress (83), and in hour hands after enforced wakefulness. Indeed, EEG SW power during NREMS is sensitive to cortical NMDA receptor function (84), increasing with NMDAR downregulation or blockade (85), in an experience-dependent manner. In rats, antagonism of NMDAR function supresses spontaneous bursting activity in cortical neurons (86), and increases EEG SW activity with a peak around 1.75 Hz (87), specifically during NREMS. A causal link between cortical activity surges in the slow-wave range and NMDAR dysregulation has also been previously observed following the application of the NMDAR-antagonist ketamine (88), as well as in models of NMDAR hypofunction, particularly after the administration of the acute chemoconvulsant pentylenetetrazole (PTZ), which enhances neuronal excitation (89). After miR-709 inhibition, downregulated genes like *Grin1,* involved in *Positive regulation of long-term synaptic depression (GO:*1900454) and *Regulation of postsynaptic membrane potential* (GO: 0060078) *in vitro, as well as Ion transport* (GO:0006811) *in vivo,* also support the notion that miR-709 inhibition may alter E/I dynamics in cortical neurons, thereby boosting δ1 wave density in an activity-dependent manner (90). Moreover, NDMAR-dependent trafficking of other ion receptors on the neuronal membrane during LTD or LTP, for activity-dependent endocytosis and reintroduction in response to excitation, is also highly reliant on a well-regulated autophagic machinery (91, 92), which we report here as perturbed after miR-709 inhibition. Altered expression of NMDAR-signalling genes, possibly as a result of dysregulated endosomal trafficking, provides a mechanistic link between the observed transcriptomic changes and the NREMS δ1 density surge after miR-709 inhibition.

Considering its target genes and their pathways, miR-709 appears to fine-tune the balance between activity-dependent plasticity processes, such as receptor trafficking and cytoskeletal remodelling, and survival of the neuron. Lower levels of miR-709, particularly in periods of prolonged wakefulness, may thus contribute to an excitotoxic environment via imbalanced autophagic/endosomal trafficking pathways (93, 94), which ultimately results in the downregulation of transcriptional levels of genes participating in glutamatergic signalling. This is also in agreement with previous observations where *Grin1* transcriptional downregulation has been observed as a response to glutamate-induced excitotoxicity (95, 96).

To conclude, the current study attributes a role for miRNAs in EEG SW regulation. Our transcriptional and electrophysiological findings underline the importance of taking developmental, temporal and spatial aspects of miRNA expression and function into consideration for future studies. Importantly, the role of autophagic and endosomal trafficking pathways in the regulation of plasticity and brain activity should be further elucidated in the context of sleep, particularly when their reciprocal interaction has been recently confirmed in several species (10, 97–99).

## Methods

### Animals and housing conditions

All mice (males; 12 weeks old) used were individually housed in polycarbonate cages (31 × 18 × 18 cm) in a sound attenuated and temperature/humidity controlled room (23-24°C, 50-60% respectively). Mice were kept under a 12h light/ 12h dark cycle (lights on at 8 AM, 70-90 lux). Animal had access to food and water *ad libitum.* All experiments were approved by the Ethical Committee of the State of Vaud Veterinary Office, Switzerland (license number 2348 and 2676), and for the conditional *Dicer* KO mice by the Ethical Committee for Animal Experimentation of the Hôpital du Sacré-Coeur de Montréal.

### Generation of conditional Dicer knockout mice

A first, non-inducible, forebrain-specific conditional *Dicer* knockout (cKO) mouse line was created by crossing homozygotes *Dicer^fl/fl^* (100) to a transgenic *α-CamKII-Cre line* (101, 102). This line was previously shown to express Cre-recombinase mainly in excitatory neurons in the forebrain from approximately 1.5 months old. The generation and characterization of the *Camk2a-Cre^+^;Dicer^fl/fl^ KO* mice was described previously (102, 103). For all experiments, *Dicer^fl/+^* mice were used as control.

### Generation of inducible conditional Dicer knockout mice

A second, forebrain-specific and inducible conditional Dicer knockout *(icKO)* mouse line was obtained by crossing hemizygote *Camk2a-Cre*ERT2 mice (104), expressing an inducible Cre-recombinase under the control of the *Camk2a* promoter *(Cre^+^;*Jackson laboratory, Bar Habor, ME, USA, stock number 012362) mice with homozygote *Dicer^fl/fl^* mice (Jackson stock number 006366). The recombination was induced in 8-weeks old animals by intraperitoneal (i.p.) injection of 1 mg tamoxifen (100 μl; Sigma-Aldrich, Buchs, Switzerland), dissolved in sunflower seed oil/ethanol (10:1, 10mg/ml final), twice a day for five consecutive days. Control animals were injected with 100 μl of sunflower seed oil/ethanol (vehicle) with the same protocol. For all the experiments, vehicle-injected *Cre^+^/Dicer^fl/fl^* and tamoxifen-treated *Cre^+^/Dicer^+/+^* mice were used as control groups. Recombination in the brain by excision of *Dicer* exon 24 was verified by PCR and qPCR, as previously described (36).

### EEG/EMGimplantation

At 10 weeks old, mice were implanted with EEG and electromyographic (EMG) electrodes under deep anesthesia (i.p.; Xylazine 10 mg/kg; Ketamine 75 mg/kg) as detailed previously (105). Briefly, six gold-plated screws (diameter 1.1 mm) were screwed bilaterally into the skull, over the frontal and parietal cortex. Two of them served as EEG recording electrodes and the remaining four as support for the electrode assembly. For recording the EMG, two gold wires were inserted into the neck muscles and served as EMG recording electrodes. The EEG and EMG electrodes were soldered to a connector and cemented to the skull. Animals were allowed to recover from surgery for approximately 5 days before they were connected to the recording cables in their home cage, and minimum of 6 days to habituate to the cables and the experimental room.

### In vivo miR-709 inhibition

In addition to the EEG/EMG electrodes, male C57BL6/J mice were implanted with a 5 mm cannula (PlasticsOne) reaching their right ventricle at 0.3 mm posterior from the bregma (AP), 0.84 mm lateral from the midline (ML), and 2.43 mm depth (DV). After habituation to the implants and the recording room, mice were injected intracerebroventricularly (ICV) by means of a Hamilton 5-μL syringe, with either LNA miR-709 specific power inhibitors (i-miR-709: 5’-CTC CTG CCT CTG CCT C-3’, Exiqon; 100mM final conc., n=7), LNA scrambled power inhibitors (scr: 5’-ACG TCT ATA CGC CCA-3’, Exiqon; 100mM final conc., n=7) or vehicle (n=6), at a speed of 0.6 μL/min. The vehicle for all injections was 4 μL of artificial cerebrospinal fluid (aCSF; composed of 150 mM Na, 3.0 mM K, 1.4 mM Ca, 0.8 mM Mg, 1.0 mM P; 155 mM Cl; pH 7.4). Post-injection, animals were left undisturbed for forty-eight hours prior to baseline recordings, as recommended by the LNA inhibitor manufacturer (Exiqon, Vedbaek, Denmark).

### Sleep recordings and data acquisition

*Dicer* cKO and ICV-injected mice were recorded at the age of 12-14 weeks, while *Dicer* icKO mice were recorded at 16 weeks. EEG and EMG signals were recorded continuously for 72 hours. During the 48 first hours, mice were left undisturbed and these two days were considered as baseline. Starting at light onset of the third day, animals were submitted to a sleep deprivation (SD) by gentle handling during 6 hours (ZT0–6). The remaining 18 hours after the end of the SD were considered as recovery. For the *Dicer* cKO mice, EEG/EMG signals were recorded using Lamont amplifiers (LaMont Medical, Madison, WI,USA) and Stellate Harmonie software (Stellate, Quebec, Canada) at a 256-Hz sampling frequency. For icKO *Dicer* and miR-709-inhibition experiments, EEG/ EMG signals were recorded using EMBLA™ hardware and Somnologica-3™ software (EMBLA, Thornton, CO,USA), and the analog signals were digitized at 2 kHz and subsequently downsampled at 200 Hz. The EEG was subjected to a discrete Fourier transformation yielding power spectra (range, 0.75–45 Hz; frequency resolution, 0.25 Hz; time resolution, consecutive 4 second epochs; window function, hamming). Activity in the 50 Hz band was discarded from further analysis because of power line artifacts in the EEG of some of the animals. Offline, the animal’s sleep-wake behavior was visually classified as Wakefulness, NREMS, or REMS based on the EEG and EMG signals as previously described (33, 36, 105). Four-second epochs containing artifacts were marked according to the state in which they occurred.

### Sleep distribution and EEG spectral analysis

Analysis of the distribution of each behavioral state was performed on 1-hour values as described previously (31). For each parameter, the two baseline days were averaged. The difference in the recovery-baseline accumulation was calculated by expressing the recovery time spent in sleep as a difference from the baseline time spent asleep for each hour. Time course analysis of EEG delta power during baseline and after SD was performed as described previously (30, 33). Briefly, the recording was divided into sections to which an equal number of 4-second epochs scored as NREMS contributed (i.e., percentiles). The baseline light periods were divided into twelve intervals; the dark periods into six. The second 6 hours of the recovery light period was divided into eight sections. In all recordings, delta power values were normalized by expressing all values relative to the mean value reached in the last 4 hours of the main rest period when delta power is minimal during baseline.

EEG spectral analysis was performed for different time periods, as described previously (30, 33). For the EEG spectra presented relative to a reference, as well as the EEG delta time course in the *in vivo* miR-709-inhibited mice, interindividual differences in overall EEG signal power were normalized by expressing EEG spectral density in each frequency bin as a percentage of baseline reference calculated as the mean total EEG power over all frequencies and behavioral states over the 48 hours of baseline. The relative contribution of the behavioral states to this individual reference value was weighted so as to avoid that, e.g., individuals that spent more time in NREMS (during which overall EEG power is higher compared to wake and REMS) obtain a higher reference power as a result.

TMT Pascal Multi-Target5 software (Framework Computers Inc., Brighton, MA, USA) was used to manage the data, Sigmastat (Systat Software Inc., Chicago, IL, USA) for statistical analyses, and SigmaPlot 12.5 (Systat Software Inc., Chicago, IL, USA) for graph generation. To assess genotype difference on different sleep-wake parameters (i.e., time spent in sleep and wake states, EEG delta power, recovery amount of sleep), one- or two-way repeated measures analysis of the variance (rANOVA) were performed. Significant effects were decomposed using *post-hoc* t-tests or Tukey’s HSD tests. Significance threshold was set to p=0.05. Results are reported as mean ±1SEM. All information regarding the statistical tests and number of animals used can be found in the respective Figure legends.

### Slow-wave parameter analysis

Raw EEG signals coming from Lamont amplifiers at 256 Hz and Embla amplifiers at 200Hz (see EEG recordings section), were imported into Matlab (v. 2020a, Mathworks Inc.) and single column vectors (uV; 200 values/sec), in addition to scored sleep-wake states from Somnologica (see above). To discriminate changes within low-frequencies of the EEG signal, raw data was filtered (Chebyshev type-II 0.5-4.5Hz, bandstop at 0.1 and 10Hz, with a zero-phase digital filtering function, filtfilt). To detect slow-waves, a custom-made Matlab algorithm was employed, based on zero-crossings and wave reconstruction closely following others, previously outlined here (33). NREMS SWs were determined through a step-by-step approach: (i) location of zero-crossings, (ii) identification of surrounding local maxima and minima, and (iii) thresholding was applied to control for SW-amplitude (see below). Data was grouped and averaged for NREMS episodes >32s. Amplitude, and period changes were expressed as a percentage of baseline ZT8-12 NREMS values, as with spectral power. These SWs were detected across baseline, SD, and recovery recordings for all NREMS episodes. Lower and upper thresholding was applied to capture bona-fide “true” NREMS SWs based on visual inspection and previously published criteria. Upper thresholds were set at 6 times the s.d. of the amplitude of all detected waveforms. The lower threshold for SW detection was calculated for each individual mouse to correct for global differences using a weighted mean rectified (Hilbert-transformed) EEG signal amplitude of all sleep/wake states during 48h of baseline (**Figures 2A and B**: Dicer^fl/+^ = 55±5 μV; Dicer^fl/fl^ = 39±4 μV; t-test, p=0.026. **Figures 2D and E**: Dicer^fl/fl^ + vehicle = 67±3 μV; Dicer^fl/fl^ + tamoxifen = 50±4 μV; t-test, p=0.003). Similarly, for the miR-709 inhibition (**Figure 4A**; aCSF = 61±3 μV; scrambled = 58±7 μV; imiR = 63±6 μV). Baseline values represent an average of two consecutive days.

### miRNA microarrays

Sixteen and 8 C57BL/6J male mice (12 weeks old) were used respectively, in experiment 1 (forebrain analysis) and 2 (cortex and hippocampus analysis). In each experiment, half of the mice were submitted to a 6-hour SD performed by the so-called “gentle handling protocol”, starting at light onset (ZT 0). The other half was left undisturbed and used as control. At the end of the SD (ZT6), animals from both groups were sacrificed and forebrain (experiment 1) or cortex and hippocampus (experiment 2) were dissected and collected. For forebrain dissection, the olfactory bulb, cerebellum, and brainstem were removed from whole brain, and the remaining part was considered as forebrain. For cortex and hippocampus dissection, two 2-mm-thick sagittal slices taken 1 mm lateral from the inter-hemispheric fissure were cut using a sagittal slicer matrix (Alto Stainless Matrices, Stoelting, CO, USA) in PBS (Life Technologies Europe, Zug, Switzerland). From the two resulting slices, hippocampi and cortex were dissected according to the Mouse Brain Atlas (106). All *s*amples were flash-frozen in liquid nitrogen and stored at −80°C until RNA extraction. Brain tissues dissected from SD and control mice were homogenized in QIAzol Lysis Reagent (Qiagen, Hombrechtikon, Switzerland). RNA was extracted and purified using the QIAGEN miRNeasy mini kit 50 (Qiagen, Hombrechtikon, Switzerland) according to the manufacturer instructions. All RNA sample amounts were measured with a NanoDrop ND-1000 spectrophotometer (ThermoFisher Scientific, Wilmington, NC, USA) and the quality of RNA samples was verified on Agilent 2100 bioanalyzer chips (Agilent technologies, Basel, Switzerland). Total RNA (100 ng) was dephosphorylated with calf intestine alkaline phosphatase (GE Healthcare Europe GmbH, Glattbrugg, Switzerland), denaturated with dimethyl sulfoxide, and labeled with pCd-Cy3 using T4 RNA ligase (GE Healthcare Europe GmbH, Glattbrugg, Switzerland). The labeled RNAs were hybridized to a Mouse miRNA Microarray (G4472B, Agilent Technologies, Basel, Switzerland) for 20 hours at 55°C with rotation. In the first experiment, a total of 16 arrays (i.e., 8 SD and 8 control animals) were used. For the second experiment, samples were pooled per group of 2 animals for each condition (SD/control) and for each brain region (cortex/hippocampus), resulting in a total of 8 arrays. After hybridization and washing, the arrays were scanned with an Agilent microarray scanner using high dynamic range settings as specified by the manufacturer. Agilent Feature Extraction Software was used to extract the data. Data were quantile-normalized and log2-transformed using the Agilent GeneSpring application. Statistical analysis was performed with the R Bioconductor package *limma* by fitting a linear model and computing moderated t-tests, comparing miRNA expression levels in the SD versus the control group. For multiple testing correction within each experiment, the Benjamini-Hochberg method was applied (107), which controls for the false discovery rate (FDR).

### In situ hybridization

A separate group of SD (n=5) and control (n=5) male mice was used to perform *in situ* hybridization on brain samples after 6 hours of SD or undisturbed sleep. Paraformaldehyde-fixed mouse brains were cryosected into 20 μm sections and hybridized with a miRCURY LNA™ microRNA Detection kit according to manufacturer’s instructions [Optimization Kit 4 *(miR-124),* Exiqon, Vedbaek, Denmark]. Briefly, sections were hybridized with a LNA miR-709 specific probe (predesigned, 5’-TCC TCCT GCC TCT GCC TCC-3’, Exiqon). A scrambled miRNA probe provided with the kit was used as negative control. The double-DIG labeled LNA miRNA detection probes were diluted at 1:625 for the scrambled, 1:1250 for miR-709 in hybridization buffer for 1 hour at 58°C. The DIG-labeled probes were detected by alkaline phosphatase-coupled anti-DIG, followed by color development (Substrate: NBT, BCIP, Roche Diagnostic AG, Rotkreuz, Switzerland).

### Primary cortical cultures

Dissociated cortical primary cultures were prepared from C57BL/6J mouse brains at embryonic day 17–18 (E17-E18; of either sex) as detailed previously (108, 109). In brief, pregnant mice (12-14 weeks old) were sacrificed by cervical dislocation, and embryos were removed and decapitated in phosphate-buffered saline at 4°C. Both cortices from 5 embryos were dissected in a solution of phosphate buffer containing HEPES, 33 mM glucose, and 40 mM sucrose. The isolated cortices were digested with a solution containing 50 U of Papain (Sigma) for 30 minutes at 37°C. Digestion was stopped by the addition of trypsin inhibitor (Sigma) for 10 minutes also at 37°C. Cells were dissociated mechanically by pipetting in Neurobasal medium (NBM) supplemented with 2% B-27 (Invitrogen), 0.5 mM Glutamax (Invitrogen), and 1% penicillin/streptomycin. After 4 triturations, the cells in suspension were transferred to a separate tube. Dissociated cells were centrifuged during 4 minutes at 150 g and re-suspended in 2 ml of complete NBM, then plated (at 1.5×10^6^ cells) in 3 ml of NBM on 35 mm dishes, precoated with 0.1 mg/ml poly-L-lysine. Cultures were maintained in a humidified CO_2_ incubator (5% CO2, 37°C) for 3 days.

### Transfection protocol

After 3 days *in vitro* (DIV3), half of the medium was removed and kept at 37°C. Subsequently, primary cortical cells were treated for 12 hours with 2 μl of Lipofectamine^®^ 2000 (Invitrogen) transfection reagent, and 2 μl of either specific miR-709 inhibitors (i-miR-709, 40 nM final conc., n=6), scrambled (scr, 40 nM final conc., n=6) or ddH2O (ctrl, n=6; for the exact constructs see Methods, *In vivo miR-709 inhibition*), diluted in a total of 100 μL Gibco™ Opti-MEM™ Reduced Serum Media (ThermoFisher Scientific), according to the manufacturer’s protocol. Forty-eight hours post-transfection, cells were lysed for RNA extraction.

### RNA-sequencing

Following treatment, RNA from primary cortical cells (DIV5) was extracted and purified using the RNeasy Mini Kit 50 (Qiagen), according to the manufacturer instructions for cells grown in a monolayer. All RNA sample amounts were measured with a NanoDrop ND-1000 spectrophotometer (ThermoFisher Scientific). RNA quality was assessed on a Fragment Analyzer (Advanced Analytical Technologies) and all RNAs had a RQN between 8.8 and 9.5. RNA-seq libraries were prepared using 1000 ng of total RNA and the Illumina TruSeq Stranded mRNA reagents (Illumina). Cluster generation was performed with the resulting libraries using the Illumina TruSeq SR Cluster Kit v4 reagents and sequenced on the Illumina HiSeq 2500 using TruSeq SBS Kit v4 reagents. The number of read counts per gene locus was summarized with *htseq-count* [v. 0.9.1,(110)] using Ensembl *Mus musculus*.GRCm38.86 gene annotation v86. Quality of the RNA-seq data alignment was assessed using *RSeQC* [v. 2.3.7,(111)]. The total number of passing filter (PF) reads was between 25-35 M, with a high alignment rate (>99%) and ~90% uniquely mapped reads on protein coding gene features. Gene body coverage profiles were deemed uniform and similar across samples and about 11.500 genes per sample were found expressed (RPKM >1). Statistical analysis was performed in R (R version 3.4.3). Genes with low counts were filtered out according to the rule of 1 count per million (cpm) in at least 1 sample. Library sizes were scaled using TMM normalization and log-transformed into counts per million or CPM [EdgeR package version 3.20.7; (112)]. Differential expression was computed with limma-trend approach (113) by fitting all samples into one linear model and adding the experimental batches in the design matrix. Moderated t-tests were used for comparing i-miR-709 injected to the scrambled group. As for the miRNA arrays, the adjusted p-values were computed by the Benjamini-Hochberg method (107), controlling for FDR.

### Weighted Gene Co-expression Network Analysis (WGCNA)

To investigate pathways impacted by miR-709 inhibition, the freely available statistical analysis software (*WGCNA* R package, (44)) was used. All 13760 genes sequenced from the inhibitor and scrambled groups (total sample n=12) were used to construct a signed weighted correlation network, by first creating a matrix of biweight mid-correlation (bicor) correlations between all pairs of genes, based on gene co-expression similarity (114). In order to create a signed network with scale-free topology (115), the resulting correlation matrix was transformed into an adjacency matrix using a power of *β*=18, according to the suggestions of the WGCNA package for n<20 samples and signed networks [(116); for scale-free topology and mean connectivity assessment on the presented data, see **Figure S4B**]. Based on the adjacency matrix, the Topological Overlap (T.O.) Matrix was calculated, incorporating the topological similarity of the co-expression network (116, 117). Average-linkage hierarchical clustering was used to group genes on the basis of the topological overlap dissimilarity measure (1-T.O.) of their network connection strengths. To identify modules in our dataset, we used the parameters *cuttreehybrid, minModulesize=200, deepsplit*=1 and a module eigengene dissimiliarity threshold *MEdissThres*=0.25. The summarized expression profile of each module was given by the corresponding first principal component (**Figure S4A)** explaining most of the variance in the dataset, defined as module eigengene (ME; **Figure S4C**). Module Significance was calculated by the Pearson correlation of each module eigengene to the treatment, and significant modules were selected based on the p-value of that correlation. The relationship of each gene to the eigengene of its clustering module, also known as module eigengene-based connectivity, or module membership, (kME) was calculated by the Pearson correlation between the expression level of the respective gene, and the module eigengene. Visualization of the interactions between the top representative kME genes for each of the four significant modules with respect to the treatment, was performed with the help of STRING v11.0 resource (45) see **Figures 5 and S5**). For the network creation, only the interaction with *highest confidence (0.9)* and *no more than 10* first shell interactors were used. Finally, the functional annotation for the green module was performed with the build-in Functional Enrichment tool of the STRING resource (45, 118), using the entire set of 13760 genes identified by RNA sequencing as background.

### Nanostring assays and analysis of gene expression

Cortical tissues dissected from i-miR-709 and scrambled-ICV-injected mice (n=5 for each group) were snap frozen and stored at −80°C. At the day of the experiment, they were homogenized in QIAzol Lysis Reagent (Qiagen), using a tissue disruptor. RNA was extracted and purified using the RNEasy Lipid Tissue Mini Kit (Qiagen) according to the manufacturer guidelines. All RNA sample amounts were measured with a NanoDrop ND-1000 spectrophotometer (ThermoFisher Scientific). RNA integrity was assessed by Qubit RNA HS Assay Kit (ThermoFisher Scientific).

Total RNA (130 ng) was added to a custom nCounter™ Elements TagSet in hybridization buffer and incubated for 20 hours at 67°C. The expression analysis was performed using a custom CodeSet designed to investigate the 107 out of the 111 transcripts that were found differentially regulated in the *in vitro* miR-709 inhibition (see **Table S3**). The samples were processed on the nCounter™ Preparation Station and the cartridge scanned at 555 fields of view (high density) on the nCounter™ Digital Analyser (Nanostring Technologies). After background removal (based on the geometric mean of negative controls), raw counts were subjected to technical normalization to the geometric mean of counts obtained for positive control probe set within each sample. After removal of one outlier sample from the i-miR-709 group, results were analysed using the nSolver 4.0 and nSolver Advanced Analysis Software v2.5 (NanoString Technologies).

Ten housekeeping genes were included in the CodeSet: *Cnot10, Aars, Lars, Asb7, Mto1, Ccdc127, Supt7l, Fam104a, Tada2b and Csnk2a2,* selected from the standard housekeeping genes that are used in nCounter™ Neuropathology panels (Nanostring Technologies). Following evaluation of housekeeping gene stability with geNorm (119), the expression levels for each probe within a sample were scaled using the geometric mean of seven out of the ten housekeeping genes *(Cnot10, Aars, Lars, Asb7, Mto1, Ccdc127, Supt7l*). The fold change of signal intensity was derived by comparing the normalized means between the i-miR-709 group and the scrambled group. The Gene Ontology (GO) pathways from the Gene Set Knowledge Base (GSKB) mouse database (46), similar to the MSigDB for human, was used for Gene Set Analysis (GSA) in the nSolver Advanced Analysis Software v2.5 (120). For the Differential Expression (DE) analysis, the scrambled (scr) group was set as reference and a simple linear regression was performed by the nSolver Advanced Analysis Software v2.5 to assess the level of expression of each gene in the i-miR-709 group (see **Figure 6A** and **B**). For the hierarchical clustering (**Figure 6C**), scores for each pathway in each sample were calculated as the first principal component of the pathway’s genes’ normalized expression, with treatment as a covariate. Pathway scores are plotted after Z-transformation.

## Supporting information

Supplemental Table S1. forebrain miRNAs-targets list

Supplemental Table S2. forebrain miRNA pathways

Supplemental Table S3. DE genes

Supplemental Table S4. Hub genes

## Acknowledgments

This study was supported by the Swiss National Science Foundation (SNF n°130825 and 136201 to PF), the state of Vaud supporting GMM, KK, and PF, and a grant from the Canadian Institutes of Health Research to VM. The *in vitro* experiments were supported by an additional grant by the Sophie Afenduli Foundation to KK. The authors acknowledge the Lausanne Genomics Technologies Facility for their vital contribution to all transcriptomic assays in this study.

