## Supplemental Figures for "Cortical miR-709 links glutamatergic signaling to NREM sleep EEG slow waves in an activity-dependent manner"

### Supplementary Figures

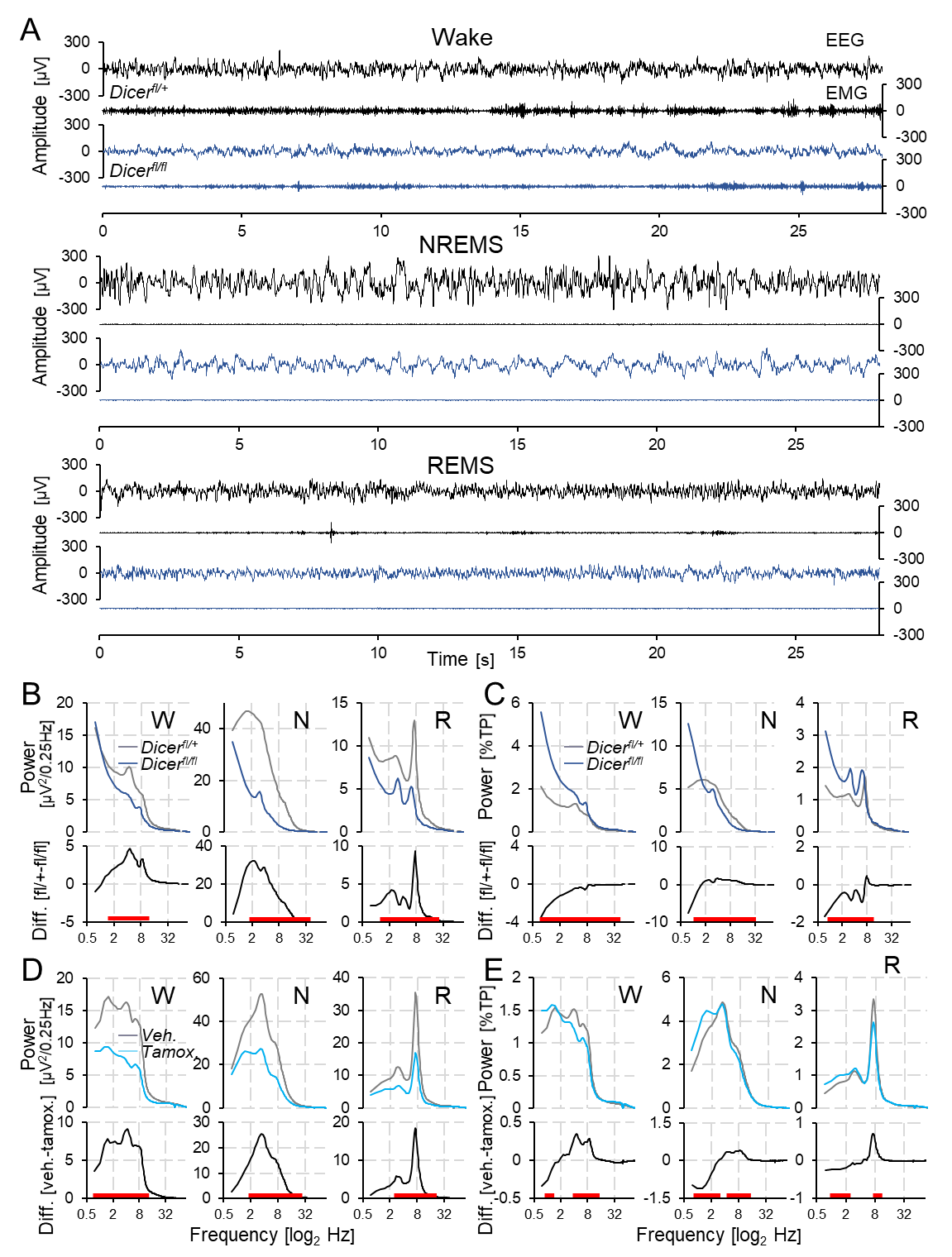

**Supplementary Figure S1: Absence of *Dicer* affects cortical EEG activity amplitude and frequency.** *Dicer*^fl/fl^ mice (dark blue) display decreased amplitude across EEG but not EMG signals compared to controls (*Dicer*^fl/+^ black), most notably during NREMS (middle), though also under waking (top) and REMS (bottom). Each example represents the first consolidated (28s of each sleep/wake state during the baseline light period), in the same two mice. A high-pass filter of 0.1Hz is applied to decrease DC offset signal artefacts. **B.** Average spectral profiles for wake (W), NREMS (N), and REMS (R), during 48h of baseline expressed in absolute values (μV2/0.25 Hz). Note the rapid decay in power from lower to higher frequencies in *Dicer*^fl/fl^ mice. Differences between group means are displayed below (black). Statistics (two-way rANOVA factors genotype × frequency); Wake: *F_167,2171_* = 3.71, p < 1E-17; NREMS: *F_167,2171_* = 22.8, p < 1E-17; REMS *F_167,2171_* = 11.9, p < 1E-17. **C.** As in B but expressed as a percentage of sleep/wake state-weighted total power (%TP) during 48h of baseline. The persistence of high power, low frequency activity (<2 Hz) in *Dicer*^fl/fl^ mice, is evident in all sleep/wake states. Statistics (two-way rANOVA factors genotype × frequency) Wake: *F_167,2301_* = 23.1, p < 1E-17; NREMS: *F_167,2171_* = 9.15, p < 1E-17; REMS *F_167,2171_* = 6.47, p < 1E-17). **D.** As in B but for inducible *Dicer* knockouts (icKO). (*Dicer*^fl/fl^ + tamoxifen; light blue) vs. controls (*Dicer*^fl/fl^ + vehicle; grey). Statistics (two-way rANOVA factors genotype × frequency; Wake: *F_167,1837_* = 11.2, p < 1E-17; NREMS: *F_167,1837_* = 14.9, p < 1E-17; REMS *F_167,1837_* = 9.14, p < 1E-17). **E.** As in C but for inducible *Dicer* knockouts (icKO). Note the presence of increased δ1 (0.75-1.75 Hz) activity during NREMS. Statistics (two-way rANOVA factors genotype × frequency; Wake: *F_167,1837_* = 3.63, p < 1E-17; NREMS: *F_167,1837_* = 6.03, p < 1E-17; REMS *F_167,1837_* = 1.88, p < 1E-17). All values represent means. Red squares represent post-hoc t-test significance (<0.05) between mice groups (B, C and D,E separately). B-D graphs are plotted on a log2 scale to emphasize lower frequency components.

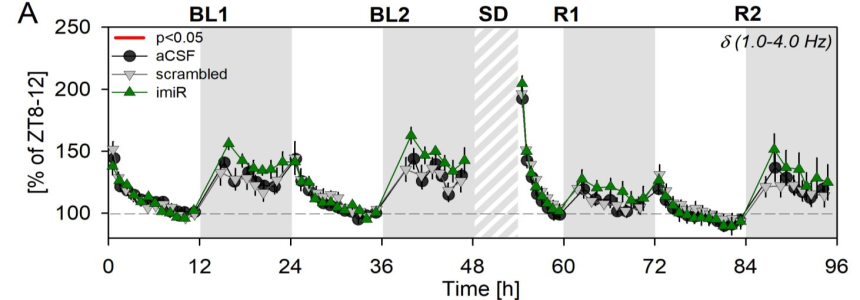

**Supplementary Figure S2: Effects of miR-709 inhibition on NREMS EEG delta power (1-4Hz).** Mean time course of NREMS EEG delta spectral power (d; 1.0-4.0 Hz) during 48h of baseline and 42h following a 6h SD at light onset. Grey areas delineate the dark periods and dashed parts the 6h SD. The power of the delta band, averaging δ1 and δ2 changes (see **Figure 4**), does not appear to be affected after miR-709 inhibition as compared to the two control groups (two way rANOVA, time p<0.001, time x condition p=0.030; post-hoc Tukey p<0.05). Note that values are given as percentage of baseline reference (ZT8-12; also see Methods).

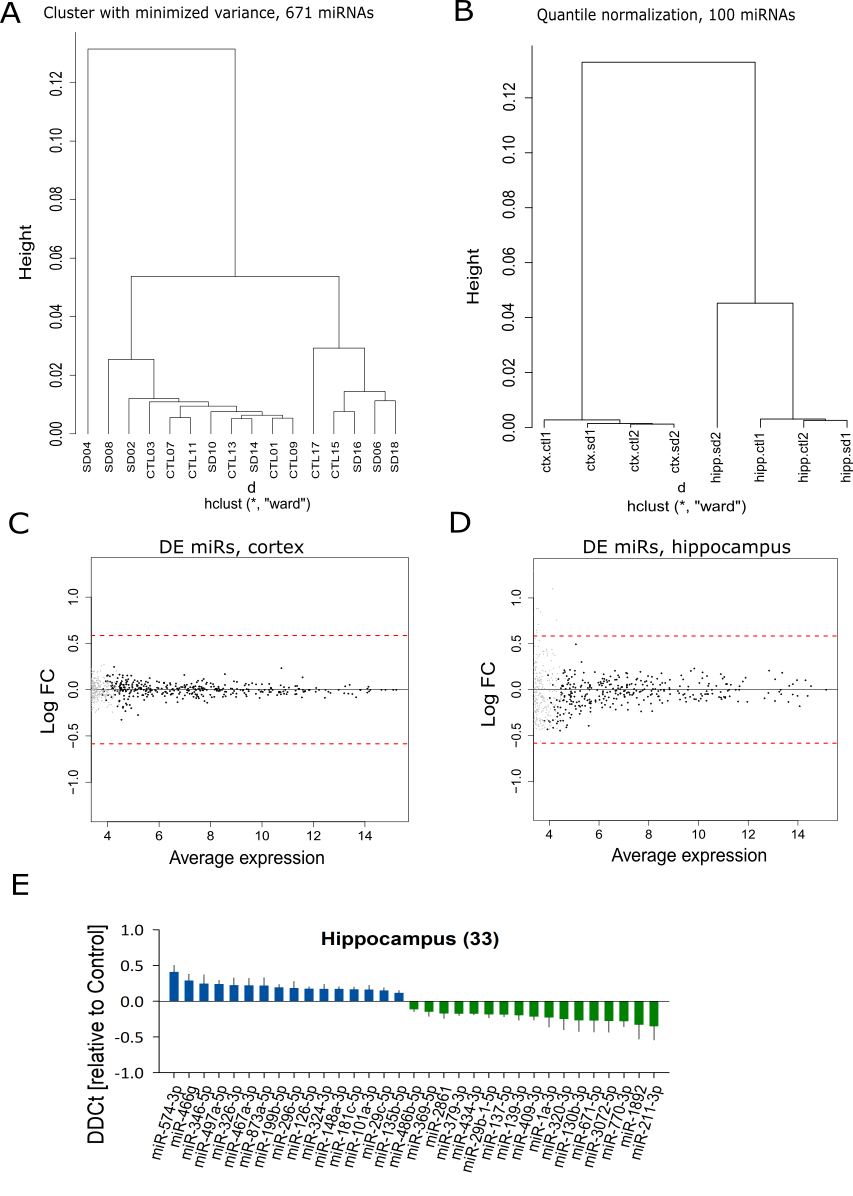

**Supplementary Figure S3: miRNA expression in the forebrain, cortex or hippocampus after 6h of sleep loss. A.** Cluster dendrogram of a total of 16 forebrain samples of both control and sleep-deprived groups, based on the expression of 671 miRNAs. Note that sample SD04 is an outlier in the clustering analysis and was therefore omitted from subsequent analysis. **B.** Cluster dendrogram of the four cortical and four hippocampal samples of the control and sleep-deprived groups, based on the expression of 1157 identified miRNAs after quantile normalization. Note that the samples cluster mainly based on the tissue, but not the condition. **C and D.** Multivariate analysis (MvA) plot of differentially expressed miRNAs in the cortex and hippocampus, respectively, of sleep deprived animals (see also **Figure 3C**). Each dot represents the differential expression of a miRNA, with grey and black dots signifying p≥0.05 and p<0.05, respectively. **E.** Mean differential miRNA expression in hippocampus after 6h SD, where 16 miRNAs increased and 17 decreased their expression (SD vs Ctrl; moderate t-statistic, p<0.05, FDR>0.05).

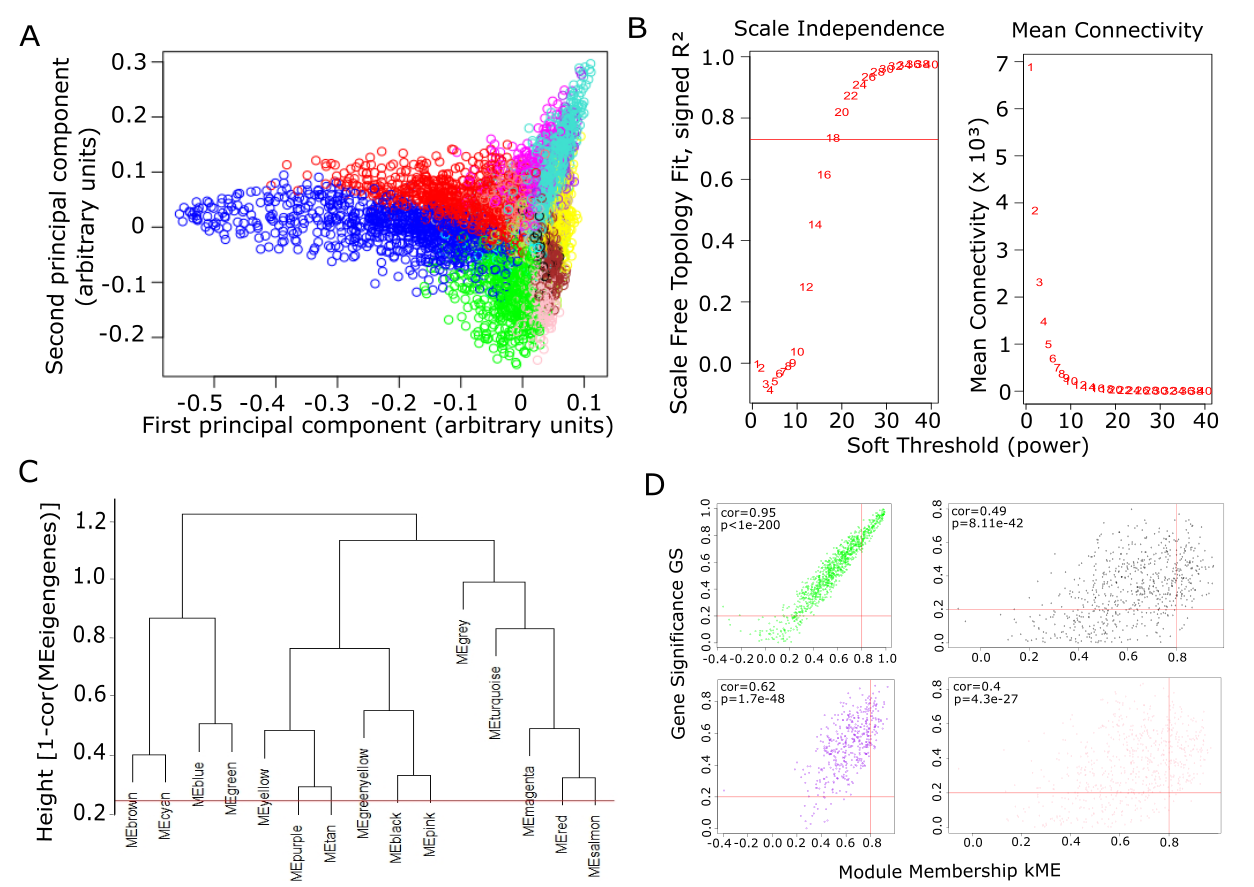

**Supplementary Figure S4. Module identification and module-module associations of the network. A.** Multidimensional Gene-network Visualization. Multidimensional scaling (MDS) plot, featuring principal coordinate analysis based on the topological dissimilarity (dissT.O.; 1-T.O.), for each module identified in the clustering analysis. The distance matrix is plotted as a function of the first two principal components. Modules that group together on this plot tend to contain genes with similar expression patterns. Note that hub genes (see **Figure 5C** and **Supp. Table S4**) for each module are located in the tip of the corresponding module. **B.** Assessment of soft-thresholding power for signed networks in WGCNA. Analysis of scale-free network topology for various soft-thresholding powers β. (Left panel) Scale-free fit index, calculated as a function of the soft-thresholding power. (Right panel) Mean connectivity in degrees, as a function of the soft-thresholding power. **C.** Hierarchical clustering of module eigengenes (MEs). Modules eigengenes summarize the modules found in the clustering analysis (also see Methods). The y-axis gives the height of dissimilarity between the module eigengenes. Branches of the dendrogram (meta-modules) group together module eigengenes that are positively correlated. No modules were merged at the selected height threshold of 0.25 (represented by the red horizontal line), corresponding to 0.75 correlation between the module eigengenes. Note that from the four significantly correlated modules (see **Figure 5B**), the black, pink and purple modules belong in the same meta-module, whereas the green module belongs to a distinct meta-module. **D.** Scatter plots for all significantly correlated modules (see **Figure 5B**), as a function of Gene Significance (GS) and Module Membership (kME) of genes. Each dot represents a gene of the respective module. Vertical and horizontal red bars depict the threshold for kME (=0.8) and GS (=0.2), respectively. Genes at the top right quadrant (GS ≥ 0.2 and kME ≥ 0.8) of each plot were identified as top hub genes (**Supp. Table S4**) for further analysis in the STRING database (see **Figure 5C)**.

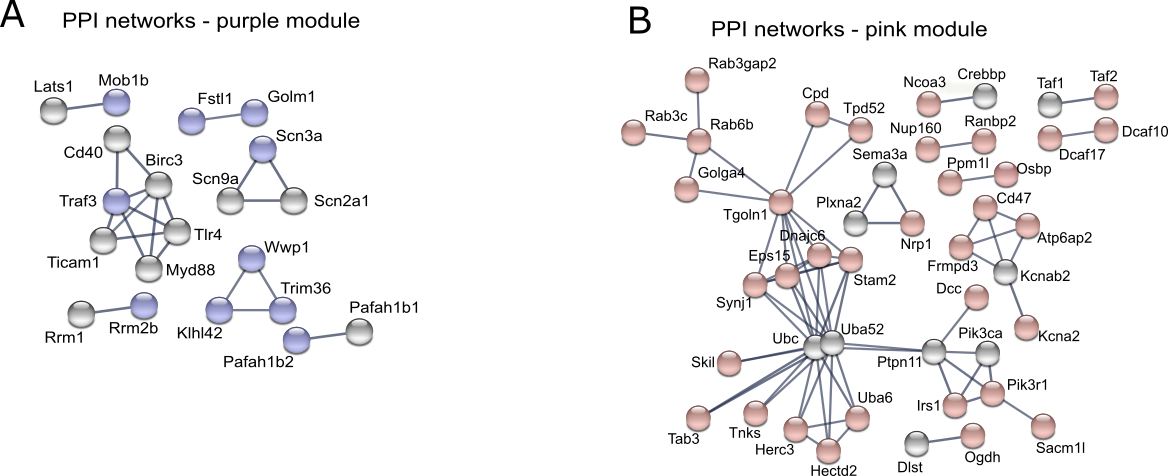

**Supplementary Figure S5. Hub genes of the modules positively correlated to the miR-709 inhibition.** Visualization of the PPI interaction networks of some of the top interconnected (hub) genes of the purple and pink modules, as calculated in STRING [only genes with high confidence (0.9) interactions are included, for the complete list of hub genes see **Supp.** **Table S4**]. Grey spheres represent genes directly linked to the green hub genes (first shell interactors; ‘no more than 10’ option), however not included in our initial dataset. Also see **Figure 5**.

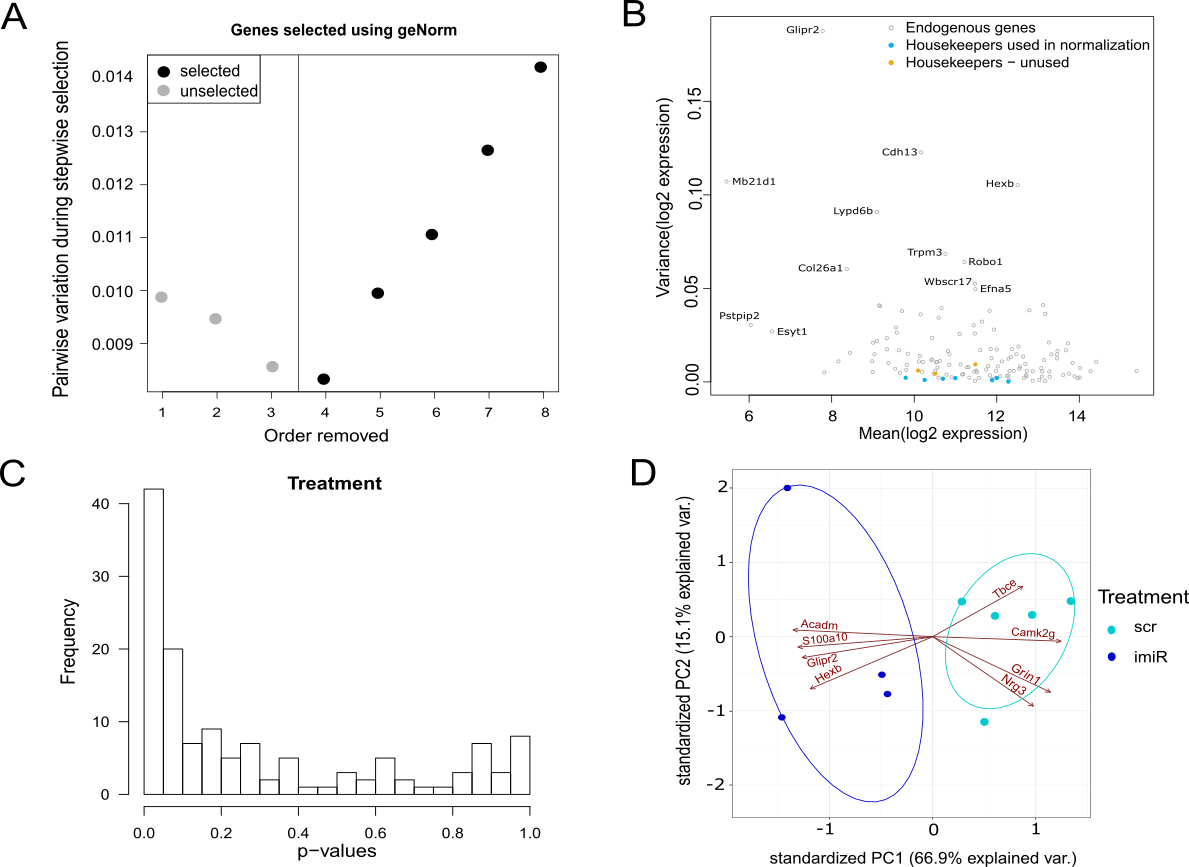

**Supplementary Figure S6. Module identification and module-module associations of the network. A.** Selection of housekeeper genes for normalization of gene expression in all samples using geNorm. Five out of the eight given housekeeping genes were selected. **B.** Scatter plot of variance as a function of mean gene expression in all samples. The selected housekeeper genes by geNorm are represented by the light blue dots at the bottom of the graph, while the three non-used housekeeping genes are of higher variance and represented by the yellow dots. All genes of interest (GOIs) are shown in grey, and only the names of GOIs with high variance or low mean expression are reported. **C.** Histogram of p-value distribution of all gene expression comparisons between treatments (inhibitor vs scrambled), providing the frequency of each p-value. The distribution is anti-conservative, suggesting a significant effect of the treatment on several genes in the dataset. **D.** PCA biplots for the common up- and downregulated GOIs between *in vitro* and *in vivo* miR-709 inhibition (see **Figure 6B**) in the Nanostring dataset. Each point in the PCA biplot corresponds to one sample. The sample’s Principal Component scores are given by the point’s coordinates. Samples with similar Principal Component scores have similar gene expression profiles and cluster together (i-miR cluster, dark blue dots; scrambled cluster, turquoise dots). The estimated region where the majority of the samples (68%) of a treatment group would be expected to fall, is represented by colored ellipses (i-miR cluster, dark blue; scrambled cluster, turquoise). Non-overlapping ellipses suggest distinctly different PC scores of treatment, as calculated from the gene expression profiles. Finally, each vector in the biplot corresponds to one gene, with the direction and length of the vector indicating its contribution to the principal component. Vectors pointing in the same direction suggest co-regulated genes.
